# PTPRJ Drives Clonal Selection in CEBPA-mutated AML

**DOI:** 10.64898/2026.07.07.736996

**Authors:** Alexandra Lubin, Catherine Hockings, Yvette Hoade, Lucy Copper, Phoebe Dace, Elizabeth Hayes, Tanvi Tambaku, Madison Hill, Amandeep Bhamra, Estere Seinkmane, Catherine Zhu, Helen Brown, Ellen Nuttall Musson, Katie-Jo Thorpe, Zhangjie Chen, Xiuyuan Chen, Silvia Surinova, Florian Grebien, Elspeth Payne

## Abstract

Transcription factor *CEBPA* is mutated in 10-15% of acute myeloid leukaemia (AML), a haematopoietic malignancy with high mortality. *CEBPA* mutations show a distinct pattern, and most patients are biallelic, carrying both an in-frame C-terminal mutation and a frameshift N-terminal mutation on opposing alleles. Rare N-terminal germline cases have 100% penetrance to AML, all with an acquired a C-terminal mutation. This suggests a selective pressure from one CEBPA mutation to develop another.

Our zebrafish models faithfully recapitulate the human disease. All biallelic mutant combinations die by 4-6 weeks of age, with pre-leukaemic haematopoietic stem cell (HSC) expansion. C-terminal and N-terminal mutants show phenotypic differences in myeloid-primed HSC and differences in the myeloid differentiation block. RNA-Seq identified differentially expressed genes in opposing vectors.

We identified phosphatase receptor *PTPRJ* as a candidate driver of clonal selection, with knock-out of *ptprja* in our fish accelerating pre-leukaemic expansion of HSC in C-terminal mutants, decelerating it in N-terminal mutants. *Cebpa*-mutant murine cells exhibit changes in differentiation and a clonal advantage with loss of *Ptprj*, which perturbs key signalling pathways.

Our data suggest that *PTPRJ* contributes to the mechanism of leukaemogenesis in CEBPA mutant AML by driving the selective pressure from each mutation to develop the other.

## Introduction

Acute myeloid leukaemia (AML) is an aggressive haematological malignancy affecting patients of all ages. The overall survival for adults with AML remains poor with less than a third of patients surviving more than 5 years^1^.

*CEBPA* is a transcription factor essential for myeloid differentiation and haematopoietic stem cell quiescence that is mutated in 10-20% of adults and children with AML. It is transcribed from a single exon and produces two in-frame protein isoforms, a shorter p30 isoform and a longer p42 isoform, with the former lacking one of the two transactivation domains. *CEBPA* mutations are known to act as founder mutations because they are observed in some families with predisposition to AML^2^. Importantly, although these families are rare, they have 100% penetrance of AML^3^. In all cases of *CEBPA*-mutated AML, additional cooperating mutations following the founder mutation are required for transformation.

Targeted therapies are already improving outcomes and treatment options for some AML patients^4^, but *CEBPA* is a transcription factor that cannot be targeted directly, particularly in germline cases where the mutation is present in all cells. Additionally, several of these therapies have failed to deliver on initial promising results. One mechanism by which success of targeted therapies is limited, or lost, is by the presence or acquisition of the cooperative oncogenic mutations. These may downregulate, or modify expression of the target, or change the cellular context in which these mutations occur^5,6^. Therefore, understanding how mutation combinations effect the disease biology, and how clonal selection of mutation combination occurs, remains crucial to the development of new therapeutics.

The most frequent secondary driver mutation in *CEBPA*-mutated AML, is the second allele of *CEBPA*. Mutations occur in a characteristic manner in the *CEBPA* gene, affecting either the N-terminal portion (*CEBPA^N-term^*) leading to production of only the shorter p30 isoform; or the C-terminal portion of CEBPA (*CEBPA^C-term^*), usually within the basic leucine zipper domain (BZIP), disrupting DNA binding. In more than 50% of cases *CEBPA*-mutated AML is biallelic, with patients harbouring both a *CEBPA^N-term^* and a *CEBPA^C-term^* mutation (*CEBPA^N-term/C-term^*)^7^. In germline *CEBPA*, the founding mutation is usually *CEBPA^N-term^*, with all individuals showing acquisition of a second hit *CEBPA^C-term^* mutation on the alternate *CEBPA* allele at the time of AML diagnosis^3,8,9^. Recurrence is common, and these patients often present with novel distinct *CEBPA^C-term^* mutations, indicating de novo leukaemia rather than relapse in many cases^8^. This highlights a strong selective pressure for this very specific biallelic mutation pattern.

Seminal murine studies have shown that *CEBPA^C-term^* mutations lead to early haematopoietic stem cell (HSC) expansion while *CEBPA^N-term^* mutations result in a differentiation block at the granulocyte-macrophage progenitor (GMP) stage^10^. These combined effects promote transformation to AML^11,12^. Interestingly, complete knockout of *CEBPA* in mice does not show a high penetrance of AML, indicating there are specific functional oncogenic features of this combination of mutated alleles^13^.

To undertake our studies, we have developed zebrafish models with *cebpa^N-term^* and *cebpa^C-term^* mutations recapitulating the mutations found in human AML. Importantly, the gene structure with multiple open reading frames controlling the expression of isoforms is conserved between zebrafish and humans^14^. *cebpa^N-term^* and *cebpa^C-term^*mutations have effects on myeloid cell development, HSC and survival. These data confirm our model faithfully recapitulates the effects of *CEBPA* mutations in mammals and gives high confidence in the relevance of our findings to human disease^15^.

Our findings indicate that the clinical prevalence of biallelic *CEBPA* mutations is not merely an additive loss of function, but rather a result of the specific, opposing, molecular phenotypes conferred by N-terminal and C-terminal mutations. We have found the *PTPRJ*, which encodes protein tyrosine phosphatase receptor J (DEP-1/CD148), is differentially expressed in opposing vectors between C-terminal and N-terminal mutations. It is known to play a key role in haematopoiesis, myeloproliferative disease, and thrombocytopenia^16–19^. Our data suggests that the interplay between *PTPRJ* and the mutational state of *CEBPA* adjusts the signalling state to push the cell toward a more favourable position within that specific *CEBPA*-mutant landscape.

## Results

### Generation of *cebpa*-mutant zebrafish models

We sought to generate zebrafish models of the *CEBPA* C-terminal and N-terminal mutations commonly observed in human AML. TALENs were designed to target the N-terminal region of zebrafish *cebpa* between the start site and 1^st^ internal reading frame, the region of human *CEBPA^N-term^* mutations in both sporadic and familial AML^20^ (Figure 1a). We identified a 5bp deletion within the TALEN cleavage site, leading to a frameshift mutation at K75, recapitulating those found in human AML, herein referred to as *cebpa^N-term^* (Figure 1b).

**FIGURE 1:**
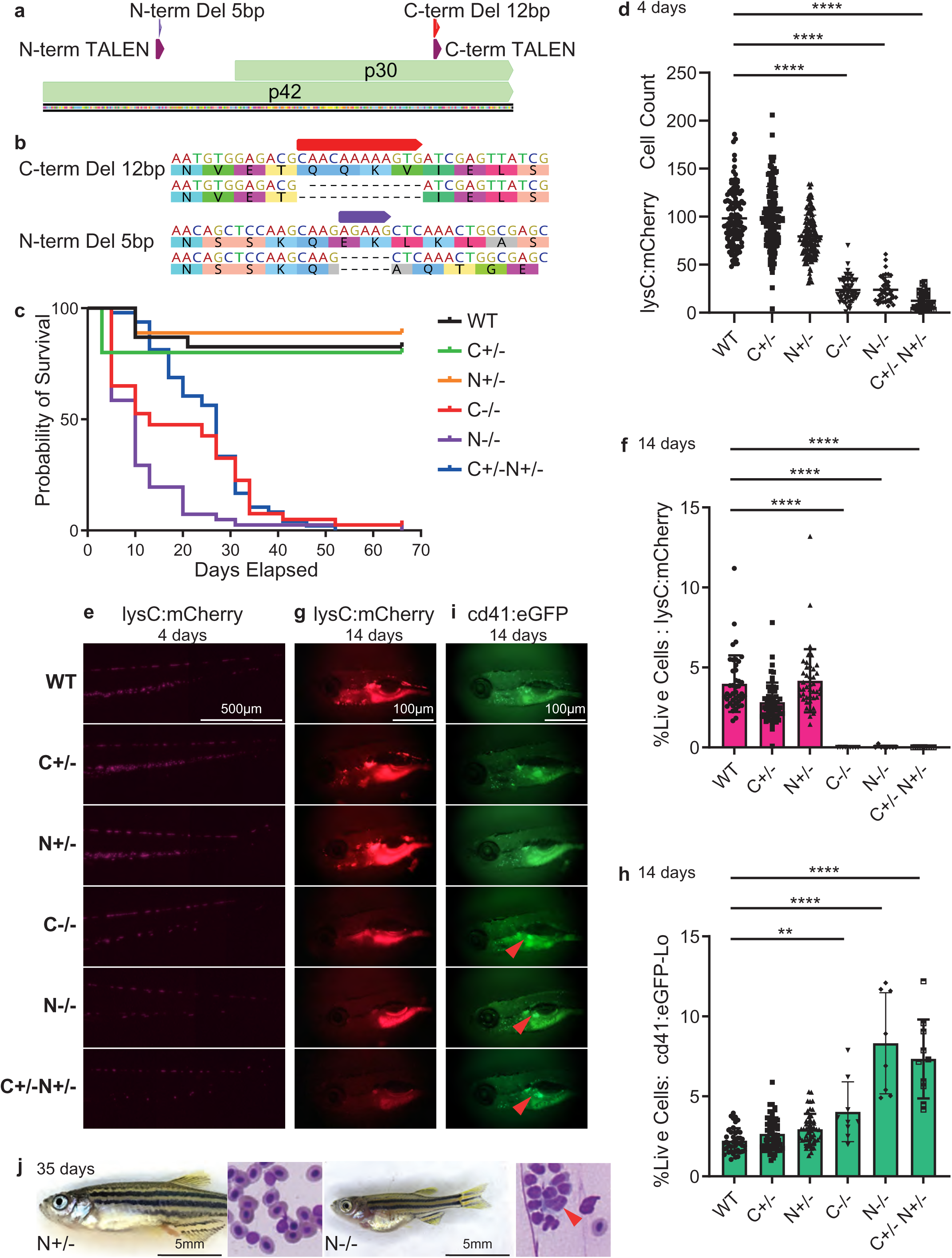
**a.** The two isoforms of *CEBPA* are well conserved between human and zebrafish, and the location of our TALENS designed to target common mutation sites. **b.** The 5bp N-terminal frameshift mutation *cebpa^N-term^* and the 12bp in-frame C-terminal deletion, *cebpa^C-term^* used in our models. **c.** Survival curve showing all *cebpa* genotypes monitored over 66 days. (N+/− = *cebpa^N-term/+^*, N−/− = *cebpa^N-term/N-term^*etc.). All biallelic mutants die by around 6 weeks. **d.** automated cell counts and **e.** images showing *lysC:mCherry+* expression in the caudal haematopoietic tissue (CHT) at 4dpf. The biallelic mutants have an absence of mature myeloid cells. **f.** quantification by flow cytometry and g. images showing *lysC:mCherry+* expression in the fish at 14dpf. The biallelic mutants have an absence of mature myeloid cells. **h.** quantification by flow cytometry and **i.** images showing *cd41:GFP+* expression in the fish at 14dpf. The biallelic mutants have a pre-leukemic expansion of HSC in the kidney (red arrows). **j.** Blood films from cardiac puncture of a leukaemic *cebpa^N-term/N-term^* juvenile alongside a *cebpa^N-term/+^*sibling show morphological differences including leukaemic-like blast cells (red arrow). One way ANOVA with GraphPad Prism. ns = p > 0.05; * = p ≤ 0.05; ** = p ≤ 0.01; *** = p ≤ 0.001; **** = p ≤ 0.0001.

For C-terminal mutants, we targeted the region in close proximity to the region of K313dup, a recurring mutation seen in biallelic *CEBPA*-mutant AML^21^ (Figure 1a). We identified a 12bp in-frame deletion around K313, again recapitulating the mutations seen in humans, herein referred to as *cebpa^C-term^*(Figure 1b).

### Biallelic *cebpa*-mutant zebrafish have impaired survival and a myeloid differentiation block

We analysed the survival of all genotypes over a period of 66 days. All biallelic mutants (*cebpa^N-term/N-term^*, *cebpa^N-term/C-term^*, *cebpa^C-term/C-term^*) showed significantly poorer survival than heterozygotes and wild-type siblings (Figure 1c). *cebpa^N-term/N-term^* juvenile show the poorest median survival of only 10 days. The vast majority of biallelic mutants die by 6 weeks. There is a sharp decline in survival in biallelic mutants between 5-10 days, when independent feeding begins, likely due to the metabolic effects of *CEBPA*^22^. Heterozygous mutants have a standard survival to juvenile age of 80-90%, and no observed differences in survival through adulthood compared to wildtype.

*CEBPA* is already well known to be a key transcription factor in myelopoiesis, and *Lysozyme C* is a key marker of myeloid lineage cells, especially mature neutrophils and macrophages. To examine the effects of our mutants on mature myeloid cells we have used the transgenic reporter *Tg(LysC:mcherry)*^23,24^. We have found an almost complete absence of mature myeloid cells in all biallelic mutants at all ages (Figure 1d-g, Supplementary Figure 1a,b).

This differentiation block is well documented in CEBPA-mutant AML and previous biallelic AML models^11,12,25,26^, highlighting the relevance of our zebrafish model. This lack of definitive myeloid cells likely contributes to the lack of tissue regeneration and poor survival on tailclipping^27^ of biallelic mutants (Supplementary Methods). Heterozygous mutants show wildtype levels of *lysozyme C* expression.

### Biallelic *cebpa*-mutant zebrafish show signs of overt leukaemia

Previous studies have shown HSC expansion prior to leukaemia-initiating blast accumulation in CEBPA-mutant AML^11^. To see if our model recapitulates this, we used *Tg(cd41:GFP-Lo)* expression as a marker for HSC (low-GFP in HSC, high-HFP in thrombocytes)^28^ in whole kidney marrow (WKM, bone marrow equivalent), which was quantified by flow cytometry. We observed a pre-leukaemic expansion of HSC in all biallelic mutants at 14dpf (Figure 1h,i), prior to juvenile death, recapitulating the human disease.

Blood films from cardiac puncture of a leukaemic *cebpa^N-term/N-term^*juvenile alongside a heterozygous (*cebpa^N-term/+^*) sibling show morphological differences (Figure 1j). While the *cebpa^N-term/+^* juvenile blood contains normally haemoglobinised and nucleated erythrocytes, the *cebpa^N-term/N-term^* shows a large number of leukaemic-like blast cells with a high nuclear to cytoplasmic ratio, and reduced numbers of poorly haemoglobinised erythrocytes.

Additionally, these homozygous mutants show a smaller size and aberrant development.

### HSC show phenotypic differences in biallelic *cebpa*-mutants during development

Having observed a full myeloid differentiation block and dysregulated HSC expansion in juvenile zebrafish, we sought to look at HSC during development. Using *Tg(cd41:GFP)* cell counts in the caudal haematopoietic tissue (CHT, fetal liver equivalent). We observed that whilst heterozygous mutants have wildtype levels of HSC at 4dpf, biallelic mutants show varying patterns of expression: *cebpa^N-term/N-term^* have increased HSC numbers, whilst *cebpa^N-term/C-term^* and *cebpa^C-term/C-term^*have decreased HSC numbers (Figure 2a,c).

**FIGURE 2:**
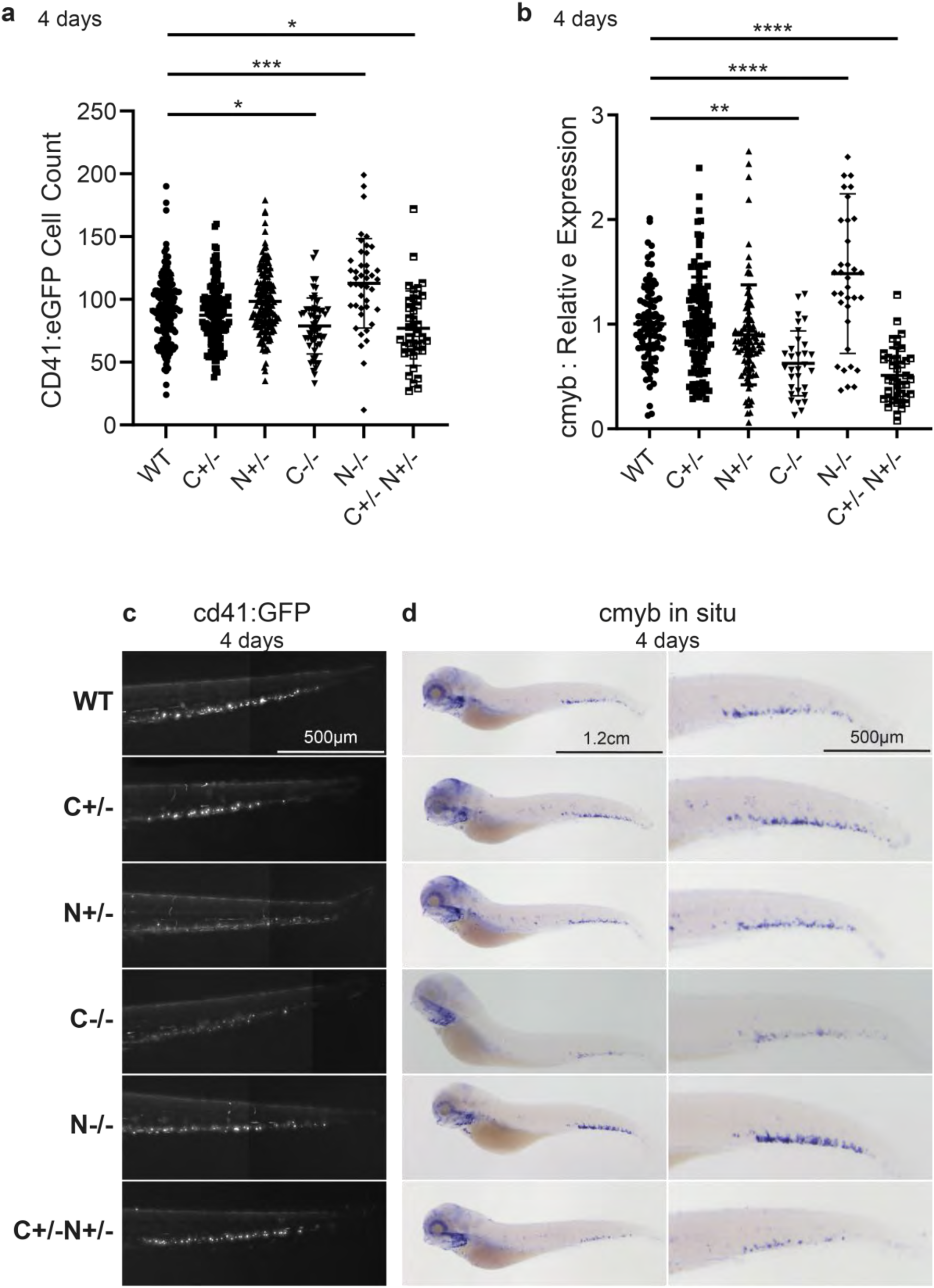
**a.** automated cell counts and **c.** images showing *cd41:GFP+* expression in the caudal haematopoietic tissue (CHT) at 4dpf. *cebpa^N-term/N-term^* have increased HSC numbers, whilst *cebpa^N-term/C-term^*and *cebpa^C-term/C-term^* have decreased HSC numbers. **b.** normalised intensity measurements by ImageJ and **d.** images showing *c-myb* expression by whole-mount in situ hybridisation in the caudal haematopoietic tissue (CHT) at 4dpf. *cebpa^N-term/N-term^* have increased HSC numbers, whilst *cebpa^N-term/C-term^* and *cebpa^C-term/C-term^* have decreased HSC numbers. One way ANOVA with GraphPad Prism. ns = p > 0.05; * = p ≤ 0.05; ** = p ≤ 0.01; *** = p ≤ 0.001; **** = p ≤ 0.0001.

We then looked at *c-myb* expression, which labels proliferating and likely myeloid primed HSC^29^. Here, the same patterns are observed but with more amplified differences (Figure 2b,d), suggesting this phenotype is driven my myeloid priming. The differences between the genotypes suggests there is a different contribution to the observed differentiation block from C-terminal and N-terminal mutations, which may contribute to clonal selection.

We also looked at *cd41* and *c-myb* expression earlier during development. During primitive haematopoiesis all biallelic mutants show increased *cd41* expression, with the largest increase in *cebpa^N-term/N-term^*, which disappears as definitive haematopoesis starts and the phenotype above takes over (Supplementary Figure 1c,d). This suggests *CEBPA* plays an important role in the regulation of HSC emergence. *c-myb* expression is largely unaffected except a small increase in expression in early *cebpa^N-term/N-term^* embryos at 32hpf, suggesting this phenotype in early development is not myeloid-driven.

### There are unique and shared gene signatures in biallelic *cebpa*-mutants

To explore these differences further we undertook bulk RNA-Seq of HSPC from each of the six genotypes. We sorted *cd41:GFP-lo* cells from zebrafish embryos and extracted RNA for sequencing (Figure 3a). The heterozygous mutants cluster with the wildtypes and show few differentially regulated genes, whilst biallelic mutants cluster independently (Figure 3b).

**FIGURE 3:**
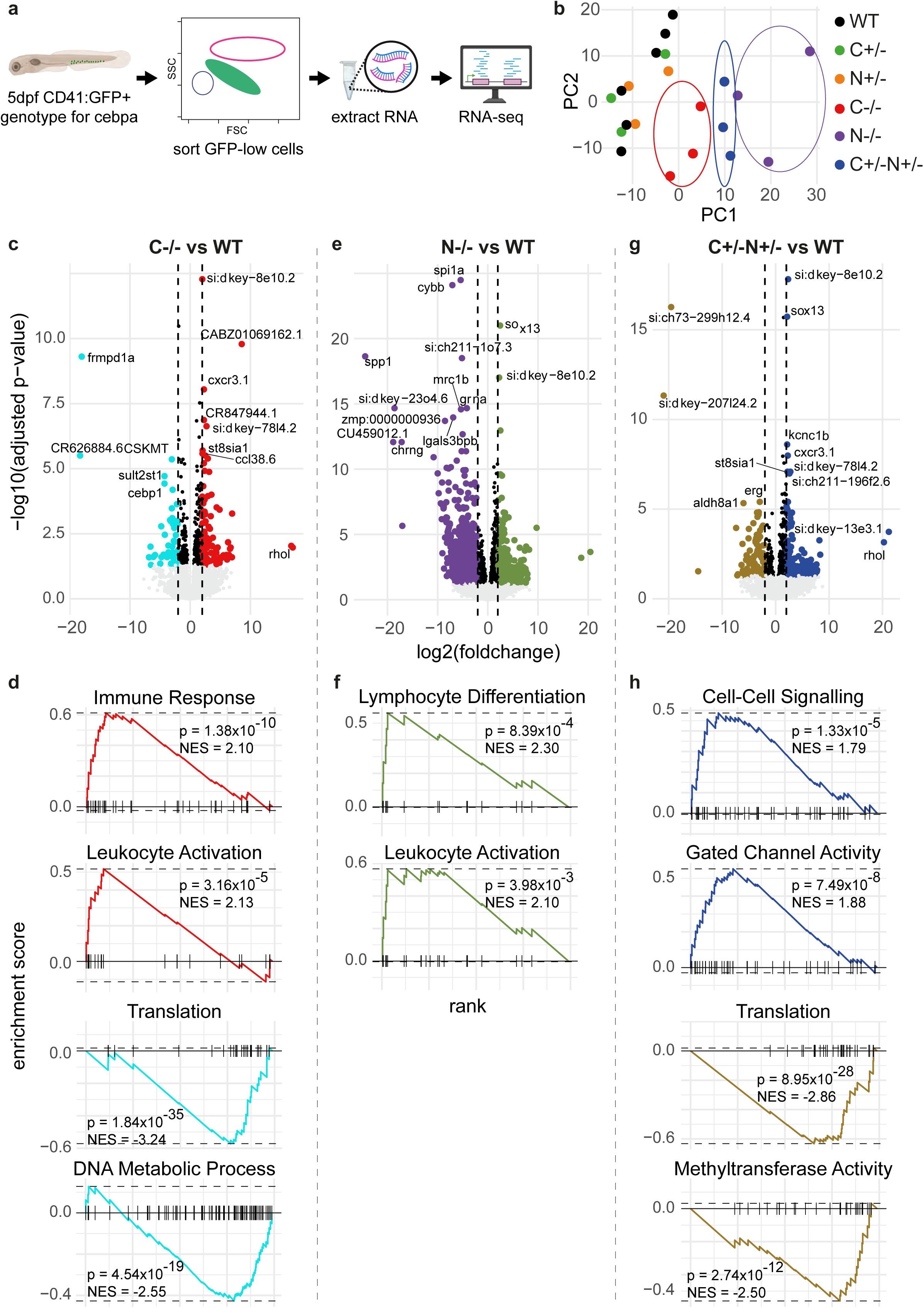
**a.** Adapted from BioRender. Payne, B. (2026) https://BioRender.com/4q1ayer. Bulk RNA-Seq on *cd41:GFP-lo+* HSC from *cebpa* mutants at 5dpf. **b.** PCA plot showing clustering of genotypes. Heterozygous mutants cluster with the wildtypes whilst biallelic mutants cluster independently. **c.** volcano plot and **d.** example of significant GSEA analysis (p<0.05) of *cebpa^C-term/C-term^* vs wildtype. **e.** volcano plot and **f.** example of significant GSEA analysis (p<0.05) of *cebpa^N-term/N-term^* vs wildtype (no significantly downregulated GSEA results). **g.** volcano plot and **h.** example of significant GSEA analysis (p<0.05) of *cebpa^N-term/C-term^* vs wildtype.

Our data revealed a large number of differentially regulated genes between each of the biallelic mutants and the wildtypes, as well as between *cebpa^N-term/N-term^* and *cebpa^C-term/C-term^*. Gene Set Enrichment Analysis (GSEA) show both unique and shared gene signatures. *cebpa^N-term/N-term^* show an increase in lymphoid activation. These mutants show increased *c-myb* expression, and *c-myb* has been shown to play a vital role in lymphoid priming and differentiation^30–32^. *cebpa^N-term/C-term^* and *cebpa^C-term/C-term^*both have a mutation in the DNA-binding domain, and show a decreased translational signature, as well as a decrease in other ribosomal biogenesis and translation pathways. The human-relevant *cebpa^N-term/C-term^* fish showed also increased cell-cell signalling.

It has been observed that null mutations in CEBPA do not cause AML, since some functional capacity enabling differentiation is required^10,13,33^. Therefore, receptors for cytokine receptors regulating differentiation – macrophage colony-stimulating factor receptors (M-CSFR), granulocyte-macrophage colony-stimulating factor (GM-CSFR), and granulocyte colony stimulating factor (G-CSFR) – were investigated. They are coded for by genes *csf1ra*, *csf2rb* and c*sf3r*, respectively, and again differences in expression are observed between genotypes (Supplementary Table 1). *csf1ra* is downregulated in *cebpa^N-term/N-term^*, *csf2rb* is upregulated in *cebpa^C-term/C-term^*, and *csf3r* is downregulated in *cebpa^N-term/N-term^* and *cebpa^N-term/C-term^*, again highlighting differences in the differentiation block and clonal selection.

We next looked at overlap between the differentially expressed genes (Figure 4a). Intriguingly, we identified 4 genes that that were differentially regulated in *cebpa^N-term/N-term^*

**FIGURE 4:**
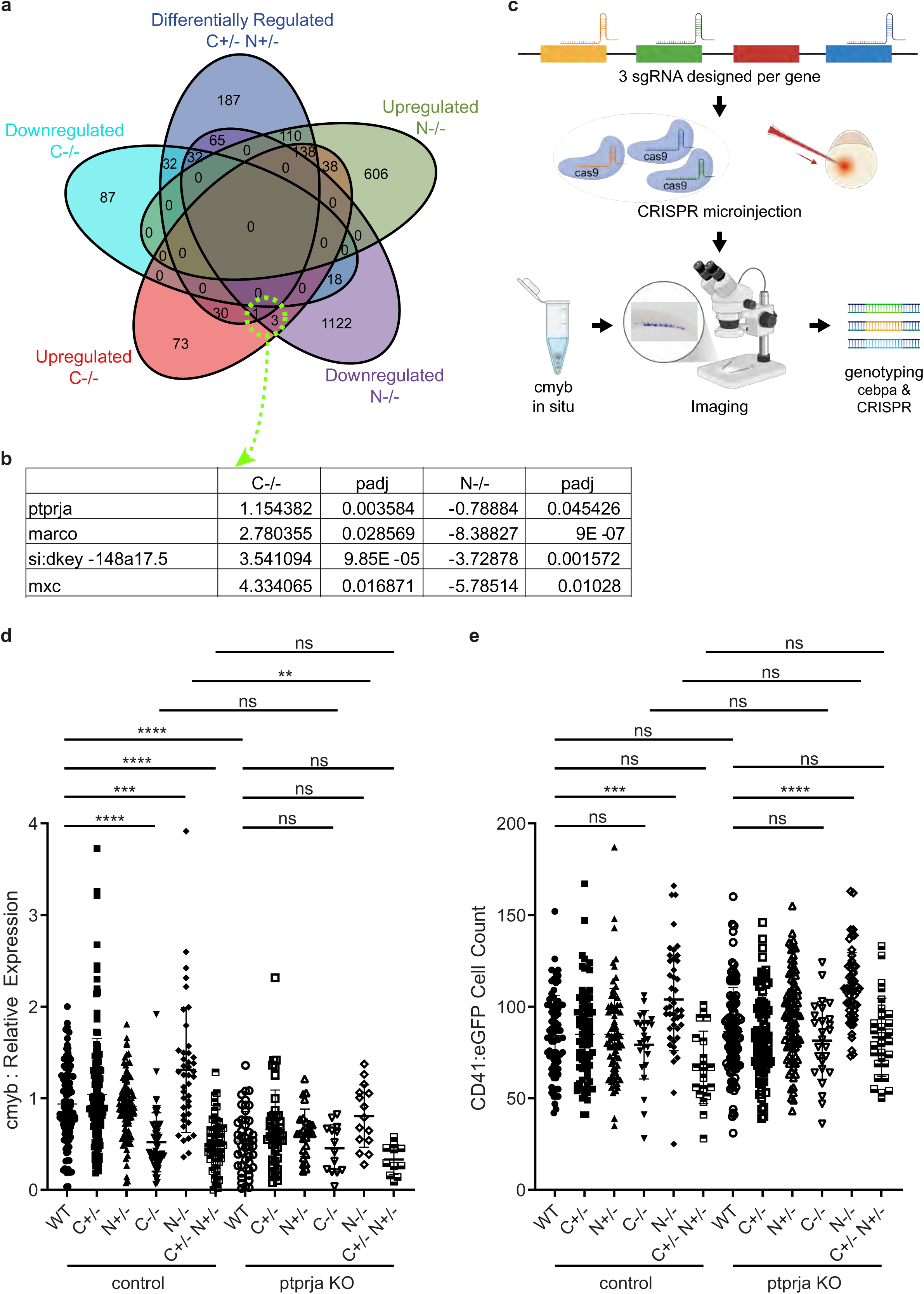
**a.** Venn diagram showing the number of upregulated and downregulated genes (p_adj_ < 0.05) in *cd41:GFP-lo+* HSC sorted from all biallelic mutant combinations vs wildtype. Up and downregulated genes are combined in *cebpa^N-term/C-term^*vs wildtype in this plot. **b.** Our analysis showed 4 genes that are upregulated in *cebpa^C-term/C-term^* but downregulated in *cebpa^N-term/N-term^*as candidates for drivers of the selective pressure to mutate the opposing allele. **c.** Created in BioRender. Payne, B. (2026) https://BioRender.com/xbequ63. CRISPR F0 knock-out screen of candidate genes using 3 sgRNA per gene. **d**. *c-myb* expression in control and *ptprja* CRISPR injected *cebpa* mutants. *ptprja* loss reduces *c-myb* expression and this is dependent on *cebpa^C-term/C-term^*. **e.** automated cell counts of *cd41:GFP+* expression in the caudal haematopoietic tissue (CHT) at 4dpf in control and *ptprja* CRISPR injected *cebpa* mutants. *ptprja* loss does not impact the counts. Two-way ANOVA with GraphPad Prism. ns = p > 0.05; * = p ≤ 0.05; ** = p ≤ 0.01; *** = p ≤ 0.001; **** = p ≤ 0.0001. and *cebpa^C-term/C-term^* HSPC, but in opposing vectors: they are upregulated in *cebpa^C-term/C-term^* and downregulated in *cebpa^N-term/N-term^*) (Figure 4a,b).

### *ptprj* is a potential novel driver of clonal selection and is dependent on functional cebpa

To investigate the role of these 4 genes we assessed effects of their loss in *cebpa* mutants using an F0 CRISPR knock-out screen (Figure 4c)^34^. An initial screen of our *c-myb* phenotype in *cebpa^C-term^* mutants identified Protein Tyrosine Phosphatase, Receptor Type J *(ptprja)* and Macrophage Receptor with Collagenous Structure *(marco)* as potential drivers, so knock-out of these was then screened across all *cebpa* genotypes (Figure 4d, Supplementary Figure 2a,b,c). This highlighted *ptprja,* as our primary candidate gene of interest. We showed that loss of *ptprja* alone and in combination with most *cebpa* genotypes reduced expression of *c-myb* compared to the control of the same genotype. However, this had no effect on expression of *c-myb* in *cebpa^C-term/C-term^* compared to controls (Figure 4d), indicating that *ptprja* reduces *c-myb*-expressing HSPC and that this is dependent on functional *cebpa* DNA binding. There were no changes in the expression of *cd41:GFP* with loss of *ptprja* (Figure 4e), showing this effect is specific to *c-myb*-expressing cells. There is also no impact on *lysC:mCherry* expression (Supplementary Figure 2d).

### Loss of *ptprja* accelerates the pre-leukaemic phenotype in *cebpa^C-term/C-term^*and alleviated in *cebpa^N-term/N-term^*

Having identified *PTPRJ* as a gene of interest we examined the impact on our leukaemic phenotype in our *cebpa*-mutant zebrafish. We used F0 knock-out of *ptprja* and screened fish at 2-8 wpf, the period biallelic mutants succumb to death (Figure 1c). We used fluorescent tumour screening and flow cytometry of whole fish or WKM to assess the leukemic phenotype. There were no changes to *lysC:mCherry* expression at any time point, with all biallelic mutants having an absence of mature myeloid cells with *ptprja* loss (Supplementary Figure 3). However, we observed differences in the expansion of *cd41:GFP* in biallelic mutants. *cebpa^N-term/N-term^* mutants have the lowest survival, with none surviving to 3 weeks or more. At 2 weeks, just as they succumb to death, there is a large expansion of *cd41:GFP* cells, which is reduced with loss of *ptprja* (Figure 1b,f). At 4 wpf, just as *cebpa^C-term/C-term^* succumb to death, we see the same expansion of *cd41:GFP* cells, but in this case this is exaggerated by loss of *ptprja* (Figure 1d,g). In the small number of *cebpa^C-term/C-term^* that survive to 8 wpf this is further exaggerated. *cebpa^N-term/C-term^* succumb to death around 3 wpf, with expansion of *cd41:GFP* cells that is unaffected by *ptprja* loss.

We also undertook proteomics on sorted HSC from *cebpa^N-term/N-term^*and wildtype siblings at 4dpf, with and without loss of *ptprja*. The low cell numbers and limited annotations in zebrafish limited this approach and the number of proteins identified. However we were able to identify a number of differentially expressed proteins when comparing *cebpa^N-term/N-term^* and wildtype with *ptprja* knock-out (Figure 5h). To elucidate mechanisms driven by *ptprja* loss in the *cebpa^N-term/N-term^*mutant background, rather than just driven by the N-terminal mutations, we performed GO analysis with FishEnrichr^35,36^ on 49 genes exclusively differentially expressed in *cebpa^N-term/N-term^*vs wildtype with *ptprja* loss, and not in the control comparison. This revealed significant downregulation of haematopoietic cell differentiation and regulation, as well as dicarboxylic acid metabolic pathways (Figure 5i). Additionally, there was significant downregulation of the pentose phosphate KEGG pathway (p = 0.036).

**FIGURE 5:**
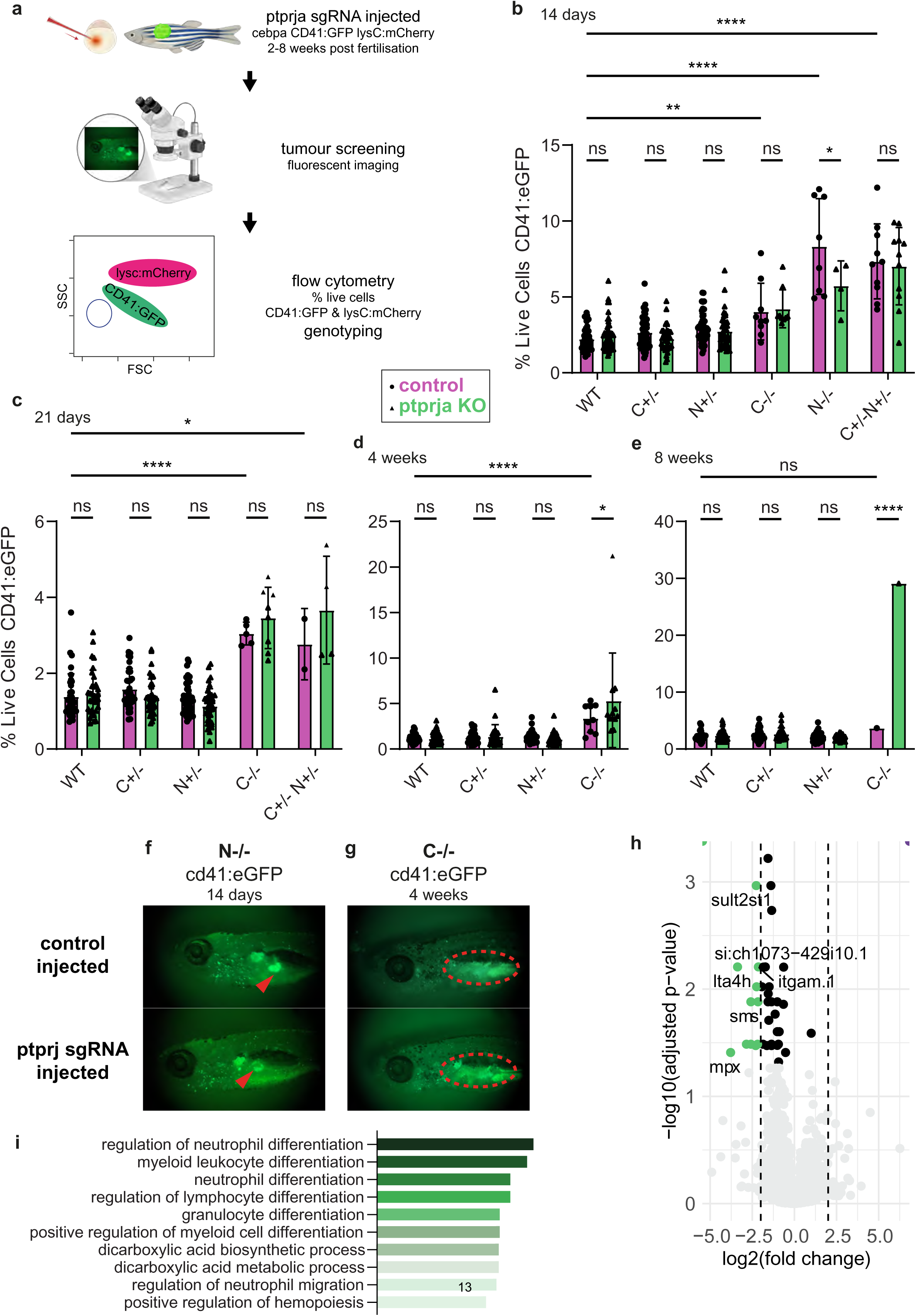
**a.** Adapted from BioRender. Payne, B. (2026) https://BioRender.com/4q1ayer. Fluorescent tumour screening and flow cytometry of whole fish or WKM to assess the leukemic phenotype. **b.** Analysis by flow cytometry of *cd41:GFP+* HSC cells in control and *ptprja* CRISPR injected *cebpa* mutants at 14 days. *ptprja* loss in *cebpa^N-term/N-term^* leads to reduced expansion of HSPCs compared with controls. **c.** Analysis by flow cytometry of *cd41:GFP+* HSC cells in control and *ptprja* CRISPR injected *cebpa* mutants at 21 days. *ptprja* loss in *cebpa^N-term/C-term^* does not change expansion of HSPCs compared with controls. **d.** Analysis by flow cytometry of *cd41:GFP+* HSC cells in control and *ptprja* CRISPR injected *cebpa* mutants at 4 weeks. *ptprja* loss in *cebpa^C-term/C-term^* increases expansion of HSPCs compared with controls. **e.** Analysis by flow cytometry of *cd41:GFP+* HSC cells in control and *ptprja* CRISPR injected *cebpa* mutants at 8 weeks. *ptprja* loss in *cebpa^C-term/C-term^* increases expansion of HSPCs compared with controls. **f.** *cebpa^N-term/N-term^* at 14 days and **g.** *cebpa^C-term/C-term^*at 4 weeks – images showing the respective decrease and increase in GFP in the kidney (red markings) with *ptprja* loss. **h.** Volcano plot of differentially expressed proteins from total proteomics of *cd41:GFP+* HSC cells at 4dpf in *cebpa^N-term/N-term^* vs wildtype with loss of *ptprja.* **i.** GO Biological Process AutoRIF analysis with FishEnrichr on 49 genes exclusively differentially expressed in *cebpa^N-term/N-term^*vs wildtype with *ptprja* loss. Two-way ANOVA with GraphPad Prism. ns = p > 0.05; * = p ≤ 0.05; ** = p ≤ 0.01; *** = p ≤ 0.001; **** = p ≤ 0.0001.

### Loss of *Ptprj* conveys a clonal advantage in *Cebpa*-mutated myeloid progenitor cells

In order to validate our findings in a mammalian model, and generate more material for proteomics, we utilised two biallelic *Cebpa*-mutant cell lines. Firstly, we used a cas9-expressing immortalised myeloid progenitor cell line derived from a mouse model of *CEBPA*-N-terminal AML that only produce the p30 isoform^12,37^, *Cebpa^N-term/N-term^* – these are non-leukaemogenic upon transplantation. Secondly, we used a cas9-expressing cell line that has the same N-terminal mutation alongside a K313dup C-terminal mutation, *Cebpa^N-term/C-term^*, derived from a tertiary transplant of an AML mouse model – these cells induce leukaemia upon transplantation^11,38^.

Viral transduction of guides targeting exon 7 of *Ptprj* resulted in an initial mutation rate of 10-20% which was monitored by NGS over 4 months. In both *Cebpa^N-term/N-term^* and *Cebpa^N-term/C-term^*cells the mutation rate grew steadily over time reaching near 100% demonstrating a positive clonal selection for *Ptprj* mutants (Figure 6a). Sequencing confirmed that in both cases these were single clones that had a clonal advantage, and both were nonsense mutations introducing a premature stop codon – a 1bp insertion and a 33bp deletion over the start of the exon respectively (Figure 6b).

**FIGURE 6:**
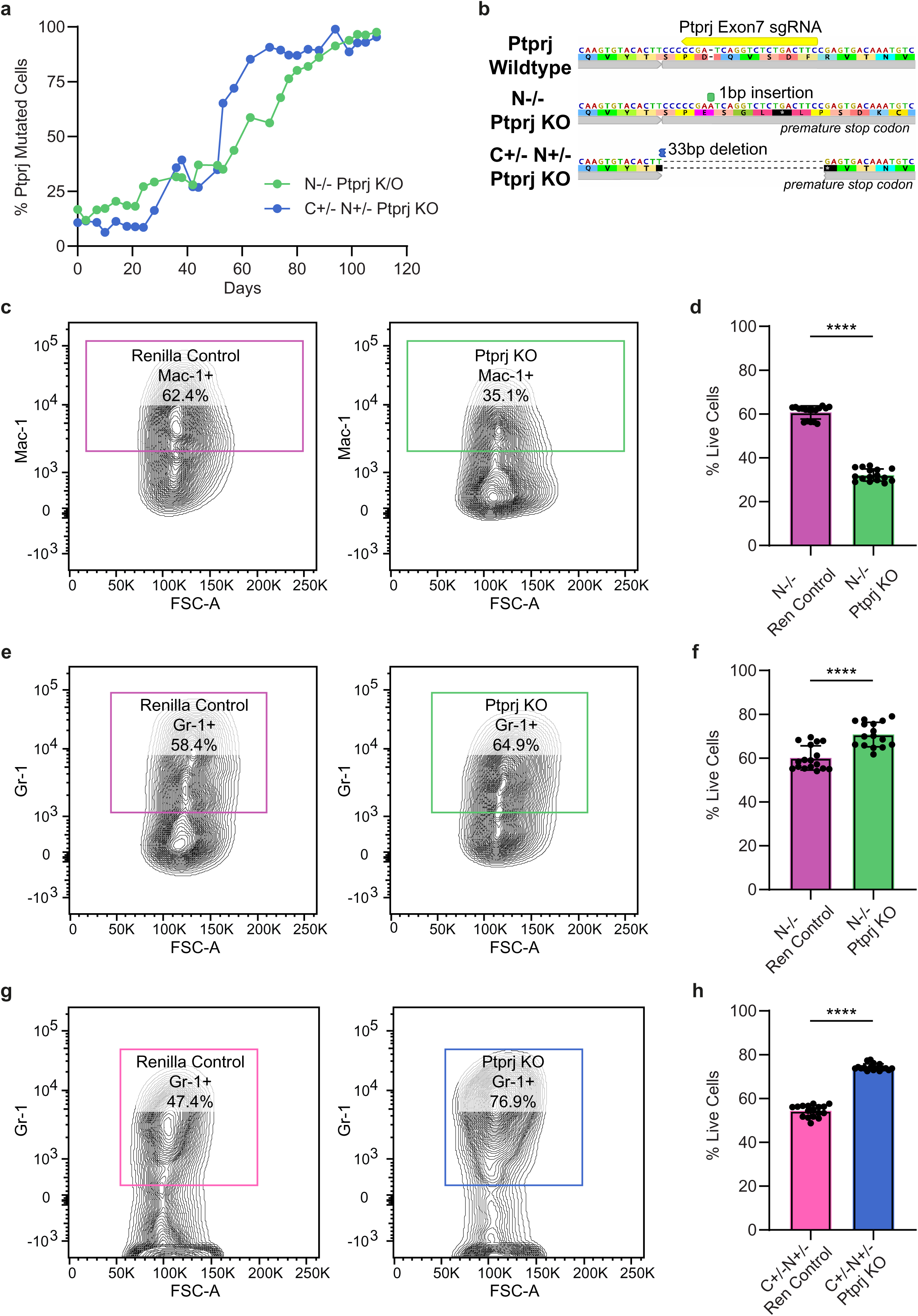
**a.** Mutation rates over 120 days post transduction of *SpCas9-Cebpa^N-term/N-term^*and *SpCas9-Cebpa^C-term/N-term^* with *Ptprj* sgRNA showing the increase in mutation rate over time. **b.** The single clones that persisted at the end of the 120 days. **c.** Example flow plots and **d.** quantification of changes in Mac-1 expression in *Cebpa^N-term/N-term^* with *Ptprja* loss showing a decrease in expression. **e.** Example flow plots and **f.** quantification of changes in Gr-1 expression in *Cebpa^N-term/N-term^* with *Ptprja* loss showing an increase in expression. **g.** Example flow plots and **h.** quantification of changes in Gr-1 expression in *Cebpa^N-term/C-term^*with *Ptprja* loss showing an increase in expression. One-way ANOVA with GraphPad Prism. ns = p > 0.05; * = p ≤ 0.05; ** = p ≤ 0.01; *** = p ≤ 0.001; **** = p ≤ 0.0001.

The *Cebpa^N-term/N-term^* cells are myeloid progenitors defined by c-Kit+;Mac-1+;Gr-1+ surface expression^12,37^. We looked for changes in these markers on loss of *Ptprj*. In the *Cebpa^N-term/N-term; Ptprj 1bp ins^* cells, there is a decrease in Mac-1 expression (Figure 6c,d), but an increase in Gr-1 expression (Figure 6e,f), compared to *Cebpa^N-term/N-term^* controls. Equally, blast-like *SpCas9-Cebpa^C-term/N-term; Ptprj -33 bp del^* cells show increased Gr-1 expression compared to controls (Figure 6g,h).

### Proliferation, stress response and DNA damage pathways are dysregulated in biallelic *Cebpa*-mutant cells with loss of *Ptprj*

To elucidate the downstream signalling cascades regulated by *PTPRJ* in the context of the common biallelic *CEBPA^N-term/C-term^* seen in patients, we performed global mass spectrometry-based phosphoproteomics comparing *Ptpr*j knockout (KO) cells against *Renilla* controls in our *Cebpa^N-term/C-term^* cells. We quantified 23,796 phosphosites across 4,166 unique proteins (Figure 7a). Utilizing a significance threshold of p < 0.05, |log2(Fold Change)| > 2 we identified 1,531 downregulated phosphosites (mapping to 601 unique proteins) and 354 upregulated phosphosites (mapping to 292 unique proteins).

**FIGURE 7:**
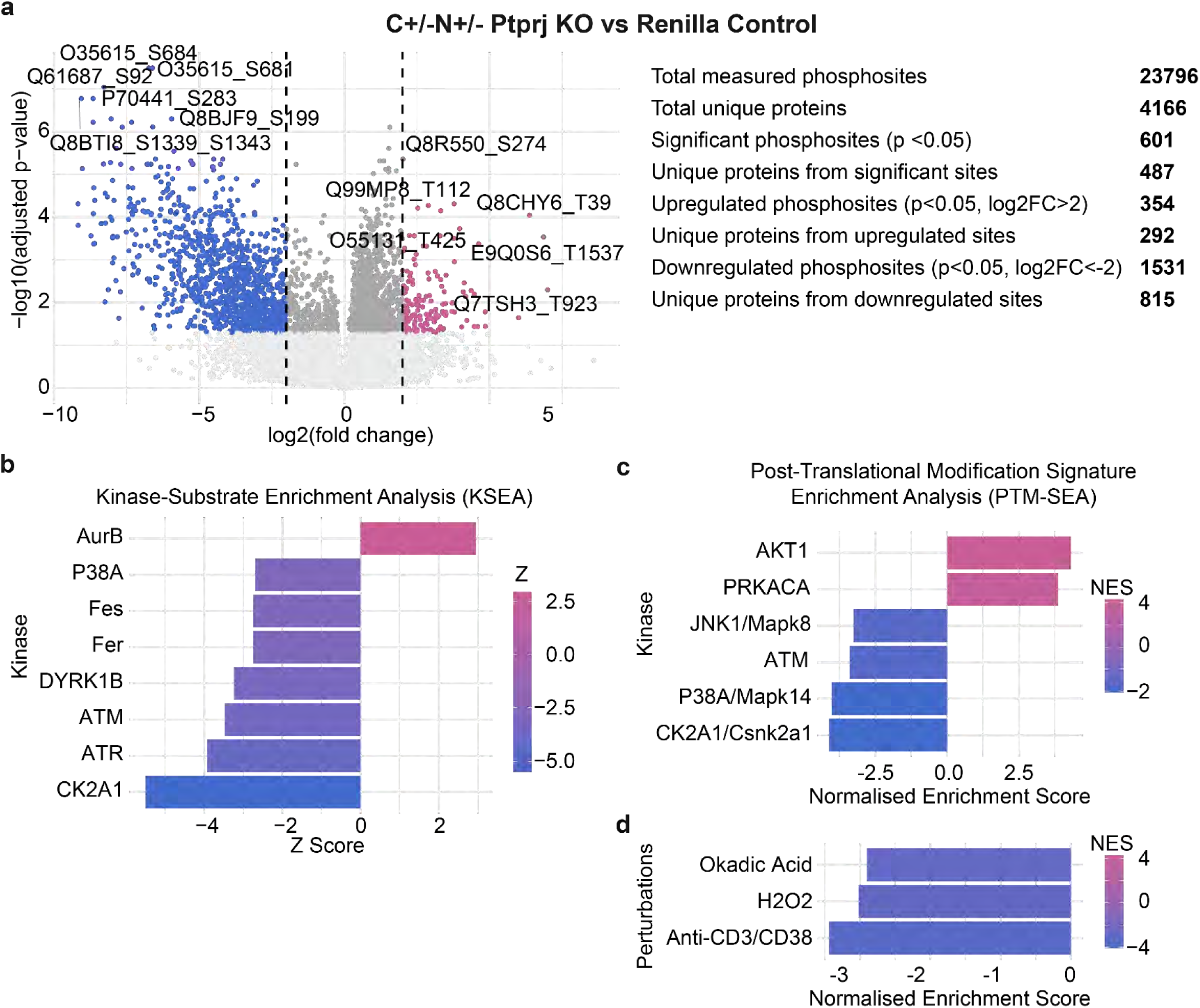
**a.** Volcano plots of significant phosphosites in *Cebpa^N-term/C-term^* cells with *Ptprj* loss compared with *Renilla* controls. **b.** Kinase Substrate Enrichment Analysis (KSEA) identification of significant kinases. **c.** Significant kinases and **d.** perturbations identified by PTM-SEA.

To identify the specific upstream enzymatic drivers responsible for this altered phosphorylation state, we conducted Kinase-Substrate Enrichment Analysis (KSEA)^39,40^ (Figure 7b). *Ptprj* loss triggered coordinated suppression of several key kinase networks. Notably, Casein Kinase 2 Subunit Alpha 1 (CK2A1) demonstrated the most pronounced reduction in substrate phosphorylation. Furthermore, we observed a marked suppression of the critical DNA damage response kinases ATR and ATM, as well as the stress-activated MAP kinase P38A. In contrast to this widespread suppression, Aurora Kinase B (AurB) showed robustly elevated activity in *Ptprj* knock-out cells.

To validate this, we also subjected the dataset to Post-Translational Modification Signature Enrichment Analysis (PTM-SEA) ^41^. PTM-SEA independently confirmed the suppression of CK2A1/Csnk2a1, P38A/Mapk14, and ATM signatures, while revealing a concurrent downregulation of JNK1/Mapk8 signalling. Conversely, *Ptprj* knock-out cells displayed enrichment for AKT1 kinase and PRKACA (Protein Kinase A) activity signatures.

Finally, we mapped these phosphoproteomic alterations against established chemical and genetic perturbation signatures. This analysis revealed that the signalling state induced by *Ptprj* loss strongly correlates with a decrease in signatures associated with Anti-CD3/CD38 stimulation, oxidative stress (H2O2) and the protein phosphatase inhibitor Okadaic Acid.

Together, these findings indicate that *PTPRJ* acts as a critical checkpoint regulating the balance between proliferation-associated cascades (AurB, AKT) and stress/DNA-damage response pathways in biallelic *Cebpa* AML models.

### Small subclones of *PTPRJ* mutant cells are present in some biallelic *CEBPA*-mutant AML samples

Finally, we wished to analyse *CEBPA^N-term/C-term^* AML patient cells to look for perturbations in *PTPRJ*. We used a custom sequencing panel to sequence the full length of *PTPRJ* and three different variant callers were used to give high-confidence mutations. From 95 patients, we identified 9 missense mutations across 8 patients, one likely germline and the remainder subclonal low variant allele frequenvy (VAF) variants (VAF < 1%) (Supplementary Figure 4a). Interestingly these clustered into just 3 single hotspot mutations: Leu24Pro, Arg425His and Ala670Thr. A search of the COSMIC database^42^, shows Ala670Thr has been detected in four cancer samples (carcinoma and glioma)^43–46^ and could impact the T-cell receptor signalling pathway. Additionally, an Arg425Cys mutation has been observed in carcinoma^47^ and could impact the same pathway. In one patient we had sequential samples from, the mutation is present in their relapse sample but not at original diagnosis.

## Discussion

Whilst a differentiation block with *CEBPA*-mutations, recapitulated in our zebrafish model, is well documented, our data reveal significant opposing differences in this block between C-terminal and N-terminal mutations. We propose that this might drive a ‘reverting’ mutation on the other allele.

Interestingly, *c-myb* hyperactivity in zebrafish has led to fish developing features of both myeloid and lymphoid malignancies^48^, and is critical for lymphoid priming and differentiation^30–32,49^. Our *cebpa^N-term/N-term^*animals with increased *c-myb* expression during development show defects in both lineages, with an absence of myeloid cells, overt features of leukaemia, and RNA-seq revealing a lymphoid activation. This signature mimics lymphoid-primed multipotent progenitors (LMPPs), consistent with previous reports that *c-myb* overexpression creates a barrier to myeloid differentiation while skewing cells toward lymphoid lineages^50^. These animals also show decreased expression of M-CSFR and G-CSFR, and *c-myb* overexpression has been shown to block differentiation by G-SCF, keeping cells in a more immature state^51^. Hence N-terminal mutations contribute to a differentiation block early in the myeloid lineage, with an expansion of immature myeloid-primed HSC and a skewing towards lymphoid-like signatures.

By contrast in *cebpa^C-term/C-term^* we observe a decrease in *c-myb* expression, and an increase in GM-CSFR. GM-CSFR in *Cebpa*-knock-out mice reactivates GMP formation, and leukemic transformation, which are blocked in *Cebpa*-null cells otherwise^52^. *c-myb* knock-out in HSC drives abhorrent but accelerated differentiation, a reduction in HSC self-renewal, and reduced multi-lineage capacity^53^. It has been shown to drive a progression of HSCs towards myeloid commitment but with defects in full maturation and a myeloproliferative state^50,54,55^. Hence C-terminal mutations contribute to a blast-like population and leukaemic transformation.

Our *cebpa^N-term/C-term^* animals, the most relevant model to human AML, show a contribution from both *cebpa^N-term/N-term^* and *cebpa^C-term/C-term^* phenotypes. This includes a balance of colony stimulating factor expression (Supplementary Table 1) and HSC expression. Our findings indicate that the clinical prevalence of biallelic CEBPA mutations is a result of the specific, opposing, molecular phenotypes conferred by N-terminal and C-terminal mutations. We propose that the *CEBPA^N-term/C-term^* combination is selected because it balances two divergent leukemic trajectories: the lymphoid-primed immature state seen in N-terminal mutants and the blast-like state seen in the C-terminal. This is substantiated by the need for some differentiation for *CEBPA*-mutant cells to be leukaemogenic^10^.

To explain these differences in differentiation, we focused on genes with opposing expression in our *cebpa* mutants (Figure 4a,b), hypothesising that these are candidate drivers of selective pressure for the common *CEBPA^N-term/C-term^*mutation combination and thus the malignant phenotype it confers.

Our data suggest that one of these genes, *PTPRJ*, acts as a critical molecular checkpoint that dictates the evolutionary trajectory of *CEBPA*-mutant cells. We have observed that perturbations in *PTPRJ* change the phenotype in biallelic *CEBPA*-mutant models. *PTPRJ*, has also recently been identified as one of the most frequently mutated genes in

*CEBPA*-mutated AML in children thereby validating the relevance of our dataset^18,56^.

Our phosphoproteomic analysis (Figure 7) reveals that *PTPRJ* is vital for regulating cell activity, with loss of *Ptprj* in our *Cebpa^N-term/C-term^*leading to a transition from a stressed, checkpoint-governed state to unregulated proliferation and a survival advantage. These cells show coordinated suppression of key stress-activated kinases which likely promotes survival and gives the cells an advantage - notably ATM (ataxia-telangiectasia mutated kinase, a master stress and DNA-damage response regulator^57^), ATR (ataxia-telangiectasia and Rad3-related kinase, which supports normal and regulated proliferation^58^) and P38α (encoded by the *MAPK14* gene, it is a key regulator of cell stress response and receptor signalling^59^). This indicates that these cells bypass traditional DNA damage response pathways that typically induce senescence or apoptosis, exaggerated by the *Ptprj* mutation.

We also see upregulation of Aurora Kinase B (AurB) phosphorylation and the simultaneously activating of this master mitotic regulator^60^, while down-regulating primary ATM and ATR, likely further allows cells to bypass replication stress checkpoints, tolerating rapid genomic duplication without initiating apoptosis or cell-cycle arrest and allowing for the expansion of pre-leukemic clones^61–63^. We observe this clonal expansion in our *Cebpa^N-term/C-term^* cells (Figure 6a).

Additionally, we see upregulation of Akt1 phosphorylation and the phosphatidylinositol 3-kinase (PI3K)/Akt pathway with loss of *Ptprj* in *Cebpa^N-term/C-term^* cells. This pathway regulates vital downstream signalling pathways including those involved in metabolism, cell proliferation, survival and growth, and hence has been identified as a therapeutic target in cancer^64–66^. It is also highly activated in AML^67,68^ and AKT1 has been specifically linked to differentiation from HSC to granulocytes^69^. This suggests that loss of *Ptprj* can push the cells into a more granulocytic progenitor state, which is exemplified by the increase in Gr-1 expression we observe in our *Cebpa^N-term/C-term^* cells with loss of *Ptprj* (Figure 6g,h).

We also see similar patterns in our *Cebpa^N-term/N-term^*cells on loss of *Ptprj* – a clonal advantage, and a shift from Mac-1 to Gr-1 suggests a skewing towards the granulocytic lineage, which is blocked by biallelic N-terminal mutations. Proteomics data in zebrafish (Figure 5i) suggests that although *ptprja* is already downregulated compared to wildtype, a complete loss still changes the phenotype. General loss of differentiation regulation with loss of *ptprja* in *cebpa^N-term/N-term^* mutants but not in a wildtype background was observed, which suggests these phenotypes are driven by the *cebpa* mutational landscape. We also see a downregulation of the pentose phosphate pathway (PPP) with *ptprja* loss, exclusively in the *cebpa^N-term/N-term^*background, which may suggest a dependency on PPP, as is common in AML,^70^ that is overcome by complete loss of *ptprja*. *Cebpa*-driven reprogramming of B-cells to myeloid lineage is directly correlated with the expression of PPP enzymes^71^. It has previously been shown that the (PI3K)/Akt pathway regulates PPP in AML^70^, and is essential for maintaining leukemic stem cells but is dispensable in normal haematopoiesis^72^. Hence loss of *ptprja* and upregulation of Akt in *cebpa* mutants may reverse this dependency.

We observed that *ptprja* loss alleviates the hyper-expansion of *cd41+* HSCs in *cebpa^N-term/N-term^*mutants. Whilst *ptprja* is already downregulated in these cells. Our data suggest that complete *ptprja* loss causes a skewing towards a more myeloid-primed, less regulated, blast-like state, which may reduce *cd41+* expression. Conversely, *ptprja* loss exacerbates the blast-like expansion in *cebpa^C-term/C-term^*mutants. *ptprja* is upregulated in these mutants and hence may be important in regulating the expansion, with its role in cell proliferation and regulation as seen in our phosphoproteomics data, leading to dysregulation on loss of *ptprja*. We also observed that the role of *ptprja* in *c-myb* expressing HSC is dependent on functional *cebpa* binding.

The N-terminal and C-terminal mutations function as two opposing vectors of clonal fitness. The N-terminal mutation drives an expansion of immature myeloid-primed HSCs through *c-myb* hyperactivity However, this priming is insufficient for full transformation. In contrast, the C-terminal mutation, by disrupting the DNA-binding domain, prevents terminal differentiation and by the downregulation of *csf1ra* and the activation of *csf2rb*, shifts the cell from a lymphoid-like state to a GM-CSFR-dependent, blast-like transformation state. *PTPRJ* acts as an accelerator of this process - fine-tuning the kinase networks to permit faster blast expansion and clonal selection of the common biallelic *CEBPA^N-term/C-term^* mutation observed in patients by balancing these opposing phenotypes to optimise for transformation. Hence these pathways provide the potential opportunity for therapeutic intervention.

## Methods

### Zebrafish Husbandry and Experimental Conditions

Zebrafish (*Danio rerio*) stocks were maintained according to standard procedures in UK Home Office approved aquaria^73^. Embryos were obtained from wild-type AB or AB/TL, or transgenic strains *Tg(itga2b:GFP)*^28^ (referred to as *Tg(cd41:GFP)*) and *Tg(LysC:mcherry)*^23^ and relevant crosses. Embryos were staged according to Kimmel *et al.*^74^ and expressed in hours/days/weeks post-fertilization (hpf/dpf/wpf). All procedures complied with Home Office guidelines.

### Generation of *cebpa* Mutants

TALEN were designed to target the N-terminal and the C-terminal regions of zebrafish *cebpa*. Wild type embryos at the 1 cell stage were injected with L and R TALEN pairs at a concentration of 300 ng/µL. Mutations were identified via PCR amplification with Phusion™ Hot Start II (Thermo Fisher) of the target loci followed by Miseq™ (Illumina) in F0 and F1 fish. STable lines were generated from a 5bp deletion within the N-terminal TALEN cleavage site *cebpa^K75fs^* referred to as *cebpa^N-term^*, and a 12bp deletion in the C-terminal TALEN site *cebpa^Q311_V314del^* referred to as *cebpa^C-term^*.

### Flow cytometry of whole kidney marrows (WKM) and individual fish

Individual fish and adult zebrafish kidneys were analysed by flow cytometry as described previously^75^. For marrow analysis, WKM were dissected from euthanised animals and suspended in ice cold Dulbecco’s phosphate-buffered saline (DPBS) with 2% fetal bovine serum (FBS) and TURBODNase 1:1000. For animals 4wpf and less, individual fish were suspended in ice cold DPBS with 2% FBS. Subsequently WKM or fish were dissociated using the gentleMACS octo dissociator (Miltenyi Biotec), passed through a 40-µM filter, and washed once with cold DPBS containing 2% FBS. Cells were resuspended in DPBS with 2% FBS and data were acquired using a BD Fortessa (Beckton Dickenson). Dead cells were excluded using 1µL Hoesct 33342 (Invitrogen) added immediately prior to analysis. Data were analysed with FlowJo Version 10.8.1 (BD Biosciences) and Prism 10 (GraphPad) software using a one-way or two-way ANOVA test.

### RNA-Seq

**Cell Sorting:** *cebpa*-mutant embryos were first genotyped at 4dpf (see Supplementary methods) and pooled per condition. At 5dpf 1000-4000 *cd41:GFP^lo^* cells were sorted directly into 500 µL Trizol (Invitrogen) via FACS using BD FACS Aria Fusion/III (BD Biosciences, USA) with a 100 µm nozzle, and stored at −20°C.

**RNA Extraction:** For RNA extraction samples were thawed on ice followed by phase separation. 100 µL of chloroform was added, samples were vortexed and incubated at toom temperature for 5 minutes. Samples were centrifuged at 12,000 × *g* for 15 min at 4°C. The aqueous phase (∼300 µL) was carefully transferred to a fresh tube. RNA was precipitated by adding 1.34 µL of GlycoBlue and 250 µL of isopropanol, a 10 min room temperature incubation and centrifugation at 12,000 × *g* for 10 min at 4°C. The resulting supernatant was removed stepwise using progressively smaller pipette tips with intermediate brief centrifugation steps (12,000 × *g* for 2 min) The pellet was washed with 500 µL of 70% ethanol and centrifuged at 7,500 × *g* for 5 min at 4°C. Ethanol was removed stepwise as described above. Pellets were air-dried at room temperature and RNA was eluted in 50 µL of nuclease-free or DEPC-treated water by incubating at 55°C for 15 min. Samples were quantified and stored at –80°C.

**Library Preparation:** Following RNA extraction libraries were prepared using the Smart-seq2 protocol^76^. Inputs calculations were based on concentration and RIN values. Reverse transcription was performed using the Maxima RT mastermix (Thermo Fisher) using 1/10 Ribolock RNase inhibitor. The reaction was incubated at 50°C for 90 min. Subsequently, KAPA HiFi PCR was utilized for preamplification. Tagmentation was performed using the Nextera XT DNA sample preparation kit (Illumina). The tagmentation reaction was assembled on ice and incubated at 55°C for 5 min. Following tagmentation, the Tn5 transposase was stripped by adding NT buffer and incubating at room temperature for 5 min. Enrichment of adapter-ligated fragments was performed via PCR using Nextera PCR master mix and index primers. Libraries were quantified and pooled and sequenced on a HiSeq 2500.

**Analysis:** Data QC was conducted using FastQC^77^ and Trimmomatic^78^ and aligned to the zebrafish genome (GRCz11) using HISAT2^79^. Differential gene expression was analysed using DESeq2 and enhanced volcano, pathway enrichment was performed using Metascape^80^ and GSEA^81^. For GSEA, pathways with zebrafish gene annotations were obtained from GO2MSIG^82^.

### F0 CRISPR Knock-Out in Zebrafish

To achieve high-efficiency biallelic disruption of our genes of interest, we employed a multi-locus CRISPR-Cas9 approach^34^ using the Alt-R™ CRISPR-Cas9 System (Integrated DNA Technologies). Three synthetic gRNAs targeting separate exons of the gene were designed using CHOPCHOP^83^. RNP complexes were prepared by incubating equal volumes of each synthetic crRNA (200 µM), tracrRNA and duplex buffer (200 µM) at 95°C for 5 min, followed by complexing this gRNA with equal volumes of Cas9 protein (61 µM) at 37°C for 5 min before pooling. Approximately 1–2 nL of the RNP mix was microinjected into the yolk of one-cell stage embryos. Mutagenesis efficiency was validated via Miseq™ (Illumina), ensuring >90% biallelic conversion using amliCan^84^ prior to phenotypic analysis.

### Tumour Screening

Fish were visualized via fluorescence microscopy using a Leica M-series stereo microscope body (Leica Microsystems) mounted on a Leica MSV269 focus column with an LabCam Ultra microscope adapter and an iPhone 13 pro. Fluorescence excitation and emission filtering (for green and red channels) was achieved using an attached Optika LED epi-fluorescence illuminator system (Optika) and images were taken with both green (for *Tg(cd41:GFP)*) and red (for *Tg(lysC:mCherry)*) fluorescent channels.

### *Ptprj* Competition Assays

Following transduction, *SpCas9-Cebpa^N-term/N-term^* and *SpCas9-Cebpa^C-term/N-term^* transduced with *Ptprj* sgRNA were monitored for mutation rate. Genomic DNA was isolated from cell samples twice weekly for 120 days. Cell suspensions (0.2 mL) were collected and centrifuged at 12,000 × *g* for 5 min to pellet the cells. Following the removal of the supernatant, the cell pellets were resuspended in 20 µL of QuickExtract DNA Extraction solution (LGC). The samples were vortexed thoroughly to ensure complete resuspension and subjected to a two-step incubation: 68°C for 15 min to facilitate cellular lysis, followed by 95°C for 10 min to heat-inactivate the extraction enzymes. The resulting genomic DNA was stored at –80°C until further experimental use. The mutation rate was analysed using Miseq™ as above. Cell lines *SpCas9-Cebpa^N-term/N-term; Ptprj 1bp ins^* and *SpCas9-Cebpa^C-term/N-term; Ptprj -33 bp del^* and *Renilla* controls were established.

### Immunophenotyping of transduced cells

SpCas9-Cebpa*_N-term/N-term; Ptprj 1bp ins_* and SpCas9-Cebpa*_C-term/N-term; Ptprj -33 bp del_* and Renilla control cell lines were aliquoted into FACS tubes (1 mL per tube), washed with 1 mL of wash buffer (DPBS with 0.1% sodium azide) and centrifuged at 300 × *g* for 5 min. Following the removal of the supernatant, cells were resuspended in 100 µL of either staining buffer (DPBS with 0.1% sodium azide and 2% FBS) or the antibody cocktail. The antibody cocktail consisted of staining buffer supplemented with anti-c-kit (anti-CD117, clone 2B8, APC, 1:400 dilution, BD Biosciences), anti-Mac-1 (anti-CD11b, clone M1/70, PerCP-eFluor™ 710, 1:200 dilution, Thermo Fisher Scientific), and anti-Gr-1 (anti-Ly-6G/Ly-6C, clone RB6-8C5, PE-Cy7, 1:200 dilution, BD Biosciences). Single colour controls were also utilised for compensation. Samples were incubated in the dark on ice for 1 hour. Following incubation, cells underwent two additional wash cycles (centrifugation at 300 × *g* for 5 min with 1 mL wash buffer). Finally, cells were resuspended in 200 µL of DPBS with 2% FBS and analysed via flow cytometry.

### Phosphoproteomics

**Cell Harvesting and Protein Extraction:** Total protein was extracted from SpCas9-Cebpa*_N-term/N-term; Ptprj 1bp ins_* and SpCas9-Cebpa*_C-term/N-term; Ptprj -33 bp del_* and Renilla control cell lines grown to 70–80% confluence in 15 cm dishes. Pelleted cells were resuspended in 300 µL of ice-cold Lysis buffer, consisting of 50 mM Triethylammonium bicarbonate (TEAB), 8 M urea, and supplemented with cOmplete™ protease inhibitor cocktail (Roche), PhosSTOP phosphatase inhibitor (Roche), and 1 mM activated sodium orthovanadate. Lysates were subjected to ultrasonic disruption using a VialTweeter (Hielscher) for five cycles (45 s on/45 s off) until the solution appeared transparent. Clarified lysates were obtained by centrifugation at 17,000 × *g* for 15 min at 10°C, and supernatants were transferred to low-protein binding tubes. Protein concentration was determined via BCA protein assay (Pierce).

**Protein Reduction, Alkylation, and Proteolytic Digestion:** Lysates were reduced with 5 mM TCEP at 37°C for 20 min and subsequently alkylated with 10 mM 2-chloroacetamide in the dark at room temperature for 20 min. Proteins were then digested with Lysyl Endopeptidase (Lys-C) at a 1:100 enzyme-to-protein ratio (w/w) for 4 h at 37°C. The urea concentration was subsequently diluted to 1.5 M with 50 mM TEAB to facilitate tryptic digestion. Trypsin was added at a 1:50 enzyme-to-protein ratio (w/w), and samples were incubated at 37°C for 4 h, followed by overnight incubation at room temperature. The digestion was quenched by acidification with 10% trifluoroacetic acid (TFA) to a final concentration of 0.5% (v/v).

**Peptide Desalting:** Peptides were desalted using C18 spin columns (capacity 35–350 µg). Columns were activated with 100% acetonitrile and equilibrated with 0.1% TFA. Samples were loaded in 400 µL aliquots, with each aliquot passed through the column twice to maximize binding efficiency (centrifugation at 70 × *g*). Columns were washed six times with 200 µL of 5% acetonitrile/0.1% TFA (centrifugation at 110 × *g*). Peptides were eluted in 225 µL of 50% acetonitrile/0.1% TFA (three 75 µL volumes). A 10% aliquot of the eluate was reserved for total proteomic analysis, and the remaining material was dried in a vacuum concentrator for storage at –80°C.

**Phosphopeptide Enrichment:** Phosphopeptide enrichment was performed using MagReSyn Zr-IMAC magnetic beads. Prior to enrichment, beads were equilibrated in loading buffer (80% acetonitrile, 5% TFA, 0.1 M glycolic acid) through three successive washes on a magnetic separator. Peptide samples (300 µg) were resuspended in loading buffer, sonicated for 5 min, and incubated with the equilibrated beads for 30 min at 1400 rpm. The bead-peptide complexes were washed sequentially to remove non-phosphorylated contaminants: once with loading buffer, twice with washing buffer 1 (80% acetonitrile, 1% TFA), and twice with washing buffer 2 (10% acetonitrile, 0.2% TFA). Each wash involved vigorous shaking at 1400 rpm for 2 min. Phosphopeptides were eluted using 1% ammonium hydroxide. To stabilize the eluted phosphopeptides, the eluate was collected directly into tubes containing 10% TFA (final concentration approximately 2%), and the elution process was repeated three times. Enriched phosphopeptides were desalted using C18 spin columns (capacity 7–70 µg) as above.

**Mass spectrometry:** nanoLC-MS/MS was performed on a Orbitrap Exploris 480 coupled to an Easy-nLC 1200 (Thermo Scientific) liquid chromatograph. 40% of each sample was analysed as 10 µL injections. Peptides were loaded on a 25 cm (75 μm ID, 1.7 μm 120 Å pore size C18 resin) Generation 4 Aurora ultimate UHPLC packed emitter column (Ion Opticks) housed in a Nanospray Flex Ion Source modified to include a column oven (Sonation GmbH) set to 35°C. Peptides were separated using a linear gradient from 6% to 38% buffer B (buffer A: 0.1% formic acid in water, buffer B: 80% acetonitrile/0.1% formic acid) over 120 min, at a flow rate of 250 nL/min. Peptides were ionised by electrospray ionisation using 1.8 kV applied immediately prior to the analytical column via a microtee built into the nanospray source with the ion transfer tube heated to 275°C and the RF-lens set to 45%. The raw data was acquired in data-independent (DIA) mode consisting of one MS1 survey scan followed by 28 MS2 scans of variable windows (Supplementary Table 2). The MS1 scan range was measuring precursors between 350-1200 m/z at a resolution of 120,000 at m/z 200. The Normalised AGC target was set to 300% (3e6 ions) (max. MS1 injection time set to “Auto”). MS2 scans were generated with the resolution set to 30,000 at m/z 200, and the normalised HCD collision energy set to 30%. The Scan Range Mode was set to “Define First mass” at 250 m/z and the Normalised AGC target set to 1000% (1e6 ions) (max. MS2 injection time set to “Auto”). The default peptide charge state was set to 2. Both of MS1 and MS2 spectra were recorded in a profile mode.

**Peptide identification, quantification and statistical analysis of discovery phosphoproteomics data:** Raw data were searched with Spectronaut (20.2.250922, Biognosys) with the directDIA™ strategy against the mouse SwissProt database (http://www.uniprot.org/, downloaded 23/05/2025) using default BGS PTM settings with the following exceptions. Precursor PEP cutoff was set to 0.01 and the “Best N Fragments Per Peptide” maximum value set to 25. Carbamidomethylation of cysteines was set as fixed modification, and oxidation of methionines, acetylation at protein N-termini, phosphorylation (on S, T or Y) were set as variable modifications. Enzyme specificity was set to trypsin/P with maximally 2 missed cleavages allowed. To ensure high confidence identifications, peptide-spectral matches, peptides, and proteins were filtered at a less than 1% false discovery rate (FDR). Label-free quantification was extracted at the MS2 area level. “PTM Analysis” was selected. Quantified phosphopeptide fragment-level data were exported as the MSstatsPTM standard report, with only the “No Decoy” filter applied. The report was filtered using R (version 4.5.1 run through RStudio (version 2025.05.1)) by only keeping fragments with F.PeakArea greater than 20 and the “EG.PrecursorID” column filtered to include precursors with phosphate localisation probabilities greater than 0.75 using the “EG.PTMAssayProbability” column. The data were then analysed within the model-based statistical framework MSstatsPTM^85^ (version 2.10.1). Data were log2 transformed, “equalizeMedian” normalised, and a linear mixed-effects model was fitted to the data. The group comparison function was employed to test for differential abundance between conditions. p-values were adjusted to control the FDR using the Benjamini-Hochberg procedure^86^. Phosphosites with missing values in one condition were assigned Inf or −Inf values to reflect the gain or loss of detection, respectively. All subsequent plots were produced in R (package ggplot2) using MSstats output files.

**Analysis:** Volcano plots were generated to visualize the distribution of fold changes and adjusted p-values, utilizing a threshold of |log_2(fold change)| > 2 and an adjusted p-value < 0.05 to define statistical significance. Kinase-Substrate Enrichment Analysis (KSEA)^39,40^ was performed to estimate kinase activity based on the differential phosphorylation of known substrate sites. Phosphosites identified as significantly regulated were cross-referenced with the PhosphoSitePlus (PSP) database^87^ to identify known kinase-substrate relationships. Kinase activity scores were calculated by integrating the log2-fold changes of all substrates assigned to a specific kinase, weighted by the statistical significance of each site. We employed Gene Set Enrichment Analysis for Post-Translational Modifications (PTM-SEA) using the PTMSigDB repository^41^. This approach maps phosphoproteomic data to annotated kinase-substrate signatures and perturbation-response signatures. Normalized Enrichment Scores (NES) were calculated to represent the direction and magnitude of pathway regulation.

