## Supplementary Methods Tables and Figures for "PTPRJ Drives Clonal Selection in CEBPA-mutated AML"

### Supplemental Methods, Figures and Legends

### Supplemental Methods

#### Genotyping All Mutant Alleles

To generate the full range of genotypes embryos were derived from the following crosses:

- *cebpa*$^{N\text{-}Term/+}$ x *cebpa*$^{N\text{-}Term/+}$
- *cebpa*$^{N\text{-}Term/+}$ x *cebpa*$^{C\text{-}Term/+}$
- *cebpa*$^{C\text{-}Term/+}$ x *cebpa*$^{C\text{-}Term/+}$

Where genotyping of live embryos is required *Tg(lysC:mCherry)* lines were used. Since biallelic mutant were found not to survive tailclipping, genotyping was done in two stages. At 3dpf embryos were screened for mCherry. mCherry negative embryos were presumed to carry biallelic mutations in *cebpa*. Their mCherry positive siblings were tailclipped[1] and DNA extracted for genotyping.

#### DNA Extraction and KASP Genotyping

DNA was extracted from embryos or finclips with HotSHOT base solution (KOH 25 mM, EDTA 200 µM). After incubation at 95°C for 30 minutes the solution was neutralised with 1x HotSHOT neutralising solution (40 mM Tris-HCl). To identify the *cebpa*$^{N\text{-}term}$ or *cebpa*$^{C\text{-}term}$ genotype we used the Kompetitive Allele Specific PCR genotyping assay (KASP, LGC Genomics).

| | |
|---|---|
| cebpa N-term KASP primer x | GAAGGTGACCAAGTTCATGCTCAACAGCTCCAAGCAAGA |
| cebpa N-term KASP primer y | GAAGGTCGGAGTCAACGGATTCAACAGCTCCAAGCAAGC |
| cebpa N-term KASP primer c | CGTAGTCTCCGCTCGCCAGTTT |
| cebpa C-term KASP primer x | GAAGGTGACCAAGTTCATGCTCCGCCGATAACTCGATCA |
| cebpa C-term KASP primer y | GAAGGTCGGAGTCAACGGATTCCGCCGATAACTCGATCG |
| cebpa C-term KASP primer c | CGGGACAAGGCGAAAATGCGCAAT |

#### Survival Curve

Embryos at 3dpf were genotyped as above, segregated by genotype and placed in the nursery. Survival was assessed by counting remaining fish numbers twice weekly. Animals were followed for 66dpf. Any remaining mCherry negative fish were genotyped by finclip at the end of the experiment to confirm.

#### Automated Cell Counting of *Tg(cd41:GFP)* and *Tg(lysC:mCherry)* Cells

Cell counts of *cd41:GFP*+ and *lysC:mCherry*+ cells in embryos were analysed using Hermes WiScan microscope (Idea Biomedical) as described previously[2]. Briefly, at 24hpf 0.003% 1-phenyl-2-thiourea (PTU) was added to the media to inhibit pigmentation and embryos were dechorionated using pronase (50 mg/mL). At the desired timepoint, live anaesthetised embryos were mounted into individual wells of a 96-well ZF plate (Hashimoto, Japan) and spun slowly for 20 seconds. Embryos were imaged in brightfield GFP and mCherry channels using 4 stitched images in x and 5 in z. Maximal projection of GFP and mCherry and best slice projection from brightfield, were performed. GFP and mCherry cell counts in the tail of *Tg(cd41:GFP)* and *Tg(lysC:mCherry)* cells, were defined using WiSoft Athena software. Following imaging embryos were removed from the plate and genotyped to correlate expression patterns with specific genotypes. Data were analysed with Prism 10 (GraphPad) software using a one-way or two-way ANOVA test.

**Blood Sampling and May-Grunwald Giemsa (MGG) Staining**

**Blood Collection:** The target for blood sampling was by cardiac puncture. Prior to blood sampling the fish were culled in tricaine and the scales overlying the puncture site were lightly dried. Non-filamented pulled needles were cut obliquely with fine scissors under a light microscope to a diameterof approximately 0.1-0.2mm. Needles were then placed gently in Heparin dissolved in saline at 1000units/ml and left until capillary action had allowed the heparin to fill approximately 3cm of the needle. Heparin was then blown out with a pipette and needles were allowed to air dry. At time of sampling the needle was inserted at 30°-45° just ventral to the spine along the axis of the zebrafish body. Needles were then gently advanced while suction was applied. Once blood was observed within the tip of the needle no further advancement was carried out and gentle suction was applied to the attached tubing until enough blood had been collected. Suction was stopped prior to removal of the needle and blood was subsequently expelled from the needle and the sample was transferred onto a glass slide and spread for MGG staining.

**MGG:** Blood films made on polylysine slides were stained with MGG to assess for morphology. Slides were fixed in 100% methanol for 5 minutes and left to air dry standing overnight. Slides were placed in neat May-Grunwald (Sigma) solution for 5 minutes, then rinsed in water twice to remove excess stain, avoiding direct contact with adherent cellular material. Slides were then placed in Giemsa (Sigma-Aldrich) (1:10 dilution) for 15 minutes, before rinsing was repeated. Having been fully air dried, slides were mounted using Permount (Electron Microscopy Services, 17986-01) and a cover slip (VWR, 631-0137). Images were taken with Hamamatsu Nanozoomer 2.0 RS.

**Whole mount *in situ* hybridization**

**Probe Generation:** An RNA probe targeting *c-myb* was generated using one-step RT-PCR (Qiagen) using the following primers:

- *c=myb*-F 5’-CCAAGTCAGGAAAACGCCACCTCG-3’
- *c-myb*-R 5’-GCTGTTGTTTAGCGGAGTTGGGCT-3’

The product was cloned into the dual promoter vector pCRII-TOPO (Life Technologies) for transcription and digoxigenin (DIG) labelling with DIG RNA Labelling Kit (Roche).

**Embryo Collection:** Whole mount *in situ* hybridizations were performed as previously described[3]. Briefly, embryos were collected, staged, and fixed in 4% paraformaldehyde (PFA) at the desired developmental time point. For embryos older than 24 hpf, 0.003% PTU was added to the media to inhibit pigmentation. Embryos were dechorionated using pronase (50 mg/mL) as required, dehydrated in a methanol series, and stored at -20°C.

**Staining:** Prior to hybridization, embryos were rehydrated in a graded methanol/PBST series and permeabilized with Proteinase K (concentration and duration adjusted per developmental stage, (5 µg/mL for 10 min at 32–36 hpf; 20 µg/mL for 35 min at 3 dpf; 30 µg/mL for 35 min at 4 dpf)). Embryos were pre-hybridized in Hyb(-) solution (50% formamide, 5x SSC, 0.1% Tween-20) at 65°C, followed by incubation with the (DIG)-labeled antisense RNA *c-myb* in Hyb(+) solution (0.25 ng/µL, Hyb(-) supplemented with 5 mg/mL torula RNA and 50 µg/mL heparin) at 65°C. Following hybridization, embryos underwent a series of stringent washes in SSC/formamide solutions.

Detection was performed using anti-DIG Fab fragments (1:5000) followed by visualization with NBT/BCIP substrate in staining buffer (100 mM Tris pH 9.5, 100 mM NaCl, 50 mM $MgCl_2$, 0.1% Tween-20). If necessary, pigmentation was removed using a bleaching solution ($H_2O_2$/formamide/SSC).

**Analysis:** Stained embryos were imaged using a Leica M205 FA stereomicroscope using a Leica DFC310 FX camera and LAS 4.0 software. All images were processed with ImageJ via Fiji version 2.0.0-rc-69/1.53v by inverting the image, converting to 8-bit and measuring the intensity. Embryos were subsequently genotyped to correlate expression patterns with specific genotypes. Data were analysed with Prism 10 (GraphPad) software using a one-way or two-way ANOVA test.

**Low Cell Number Proteomics**

To facilitate proteomic analysis from low-input cell populations, protein extraction and digestion were performed using a modified protocol optimized for minimal loss.

**Cell Collection and Lysis:** Cell suspensions from 20 pooled 4dpf *cebpa*$^{N\text{-}term/N\text{-}term}$ and *cebpa*$^{+/+}$ embryos were prepared as above. 1000 live *cd41:GFP*+ cells isolated via FACS as above directly into low-protein-binding microcentrifuge tubes (Eppendorf) containing 20 µL

LC-MS grade water and immediately flash-frozen on dry ice before storage at -80°C until use. To ensure efficient protein denaturation, samples underwent three cycles of heating at 90°C for 5 min, followed by snap-freezing on dry ice for 5 min. Lysates were then centrifuged at 17,000 × *g* for 1 min and subjected to sonication in a water bath for 15 min to disrupt cellular structures and shear genomic DNA.

**Digestion and Desalting:** Samples were centrifuged again at 17,000 × *g* for 3 min. Proteins were digested using mass spectrometry-grade trypsin (40 ng/µL in 100 mM TEAB, pH 8–9). Based on estimated protein content of 60 ng per sample : the minimum of 100 ng trypsin was used. Digestion proceeded at 37°C for 4 h (shaking at 500 rpm), followed by an overnight incubation at room temperature. The reaction was quenched to a final concentration of 0.1% trifluoroacetic acid (TFA) and adjusted to pH 3–4.

Peptides were desalted using 0.6 µL C18 ZipTips. Tips were conditioned with 100% acetonitrile and equilibrated with 0.1% TFA. After sample loading (passed through twice to maximize binding), tips were washed with 5% acetonitrile/0.1% TFA. Peptides were eluted using 50% acetonitrile/0.1% TFA, dried in a vacuum concentrator, and stored at -20°C until analysis. Prior to LC-MS, samples were reconstituted in 1% formic acid.

**Mass spectrometry:** nanoLC-MS/MS was performed on a Orbitrap Exploris 480 coupled to an Easy-nLC 1200 (Thermo Scientific) liquid chromatograph. Eighty percent of each sample was analysed as 8 µL injections. Peptides were loaded on a 25 cm (75 µm ID, 1.7 µm 120 Å pore size C18 resin) Generation 4 Aurora ultimate UHPLC packed emitter column (Ion Opticks) housed in a Nanospray Flex Ion Source modified to include a column oven (Sonation GmbH) set to 35°C. Peptides were separated using a 2-step gradient from 4% to 27% buffer B (buffer A: 0.1% formic acid in water, buffer B: 80% acetonitrile/0.1% formic acid) over 25 min, then 27% to 50% buffer B over the next 10 min, at a flow rate of 150 nL/min. Peptides were ionised by electrospray ionisation using 1.8 kV applied immediately prior to the analytical column via a microtee built into the nanospray source with the ion transfer tube heated to 275°C and the RF-lens set to 45%. The raw data was acquired in data-independent (DIA) mode consisting of one MS1 survey scan followed by 13 MS2 scans of 41 m/z windows (0.5 m/z overlap). The MS1 scan range was measuring precursors between 400-920 m/z at a resolution of 120,000 at m/z 200. The Normalised AGC target was set to 300% (3e6 ions) (max. MS1 injection time set to “Auto”). MS2 scans were generated with the resolution set to 60,000 at m/z 200, and the normalised HCD collision energy set to 27%. The Scan Range Mode was set to “Define First mass” at 250 m/z and the Normalised AGC target set to 1000% (1e6 ions) (max. MS2 injection time set to “Auto”). The default peptide charge state was set to 2. Both of MS1 and MS2 spectra were recorded in a profile mode.

**Protein identification and relative quantification:** Raw data were searched with Spectronaut (18.7.240506, Biognosys) with the directDIA™ strategy against the zebrafish UniProtKB database (http://www.uniprot.org/, downloaded 03/09/2024, 46,559 entries) using default BGS settings with the following exceptions. Precursor PEP cutoff was set to 0.01. Oxidation of methionines and acetylation at protein N-termini were set as variable modifications (maximum of 3 variable modifications on a peptide). Enzyme specificity was set to trypsin/P with maximally 2 missed cleavages allowed. To ensure high confidence identifications, peptide-spectral matches, peptides, and proteins were filtered at a less than 1% false discovery rate (FDR). Label-free quantification was extracted at the MS2 area level. Quantified fragment-level data were exported as the MSstats standard report, with only the "No Decoy" filter applied. The report was filtered using R (version 4.5.1 run through RStudio (version 2025.05.1)) by only keeping fragments with F.PeakArea greater than 20. The data were then analysed within the model-based statistical framework MSstats[4] (version 4.16.1). Data were log2 transformed, "equalizeMedian" normalised, and a linear mixed-effects model was fitted to the data. The group comparison function was employed to test for differential abundance between conditions. p-values were adjusted to control the FDR using the Benjamini-Hochberg procedure[5]. Proteins with missing values in one condition were assigned Inf or -Inf values to reflect the gain or loss of detection, respectively. All subsequent plots were produced in R (package ggplot2) using MSstats output files.

**Data Analysis:** Data cleaning, volcano plots and statistical summaries were performed in R (v4.5.2) using the tidyverse package suite[6]. Volcano plots were generated to visualize the distribution of fold changes and adjusted p-values, utilizing a threshold of |log_2(fold change)| > 2 and an adjusted p-value < 0.05 to define statistical significance. Functional analysis of significant proteins was performed with FishEnrichr[7,8], a gene list enrichment analysis tool for zebrafish, using GO Biological Process AutoRIF analysis. Results were considered significant with p-adjusted < 0.05.

#### *Cebpa* Cell Culture Methods

*SpCas9-Cebpa*$^{N\text{-}term/N\text{-}term}$ and *SpCas9-Cebpa*$^{C\text{-}term/N\text{-}term}$ cells were a kind gift from Florian Grebien[9-12].

**Culture Media Conditions:** Cells were cultured in RPMI 1640 medium supplemented with 10% Fetal Bovine Serum (FBS), 2 mM L-glutamine, 100 U/mL penicillin, 100 µg/mL streptomycin, and 5 ng/mL mouse IL-3. For initial recovery and expansion of lines thawed from liquid nitrogen, "recovery medium" was utilized, supplemented with 50 ng/mL mouse IL-3, 20 ng/mL mouse IL-6, and mouse SCF (a gift from Florian Grebien, 1:50 dilution). All cytokine stocks were prepared in PBS containing 0.1% bovine serum albumin (BSA). Cultures were maintained in a humidified incubator at 37°C with 5% $CO_2$.

**Thawing and Expansion:** Cells stored in liquid nitrogen were rapidly thawed in a 37°C water bath, transferred to 10 mL of culture medium, and centrifuged at 300 × *g* for 5 min. Pellets were resuspended in 3 mL of recovery medium and seeded into 6-well plates. Cell viability was assessed via trypan blue exclusion and hemocytometer counting.

**Passaging:** Cultures were monitored for density and health. Depending on confluence cells were passaged by centrifugation (300 × *g* for 5 min) and resuspension in fresh media every 2-3 days.

**Lenti-X 293T Cell Culture**

**Culture Conditions:** Lenti-X 293T cells (Takara Bio) were maintained in high-glucose DMEM supplemented with 10% FBS, 4 mM L-glutamine, 100 U/mL penicillin, and 100 µg/mL streptomycin. Cultures were kept in a humidified incubator at 37°C with 5% $CO_2$.

**Thawing and Passaging:** Cryopreserved cells were rapidly thawed in a 37°C water bath, diluted in 10 mL of pre-warmed complete medium, and centrifuged at 100 × *g* for 5 min. Pellets were resuspended in 10–15 mL of fresh medium and seeded into 10 cm culture dishes.

For passaging, cells were washed with PBS and detached using trypsin-EDTA incubation (3–5 min at 37°C). Following detachment, cells were resuspended in complete medium, centrifuged at 100 × *g* for 5 min, and re-plated at appropriate densities to maintain optimal confluency.

**Cloning of sgRNA Expression Plasmids**

**sgRNA design:** sgRNA targeting *Ptptj* was designed with https://www.vbc-score.org/. 3 guides were designed and the highest ranking one, targeting exon 7, was selected following successful transfection. Controls were used as per the published protocol[10,13].

| Target | Guide (inc PAM) | F Primer | R Primer |
|---|---|---|---|
| Ptprj Exon 7 | GAAGTCAGAGACCTG ATCGGGGG | CACCGAAGTCAGAGAC CTGATCGG | AAACCCGATCAGGTC TCTGACTTC |
| Renilla Control | GGATGATAACTGGTC CGCAGNGG | CACCGATGATAACTGG TCCGCAG | AAACCTGCGGACCAG TTATCATC |
| Rpa3 Control | GCTGGCGTTGACGCG CGCTTNGG | CACCGCTGGCGTTGA CGCGCGCTT | AAACAAGCGCGCGTC AACGCCAGC |

**Oligonucleotide Annealing and Phosphorylation:** Complementary forward and reverse sgRNA oligonucleotides (IDT) were phosphorylated and annealed using T4 Polynucleotide Kinase (PNK) in 1x T4 Ligation Buffer with ATP. The reaction mixture (10 µM final concentration per oligo) was incubated at 37°C for 30 min, heated to 95°C for 5 min, and then cooled to 25°C at a rate of 0.1°C/sec. The annealed products were diluted 1:200 in nuclease-free water prior to ligation.

**Ligation and Transformation:** The linearized LentiGuide-Puro-P2A-EGFP backbone (addgene 137729) was ligated with the annealed sgRNA inserts at a 1:10 molar ratio using T4 DNA Ligase. Ligation reactions were incubated at room temperature for 20 min before transformation into *Stbl3* chemically competent *E. coli*. Bacteria were recovered in SOC medium for 1 h at 37°C and seeded onto LB-agar plates supplemented with carbenicillin (50 µg/mL).

**Plasmid Purification:** Individual colonies were picked and cultured in LB broth containing carbenicillin. Clones were expanded into 30–40 mL LB cultures for large-scale plasmid purification using the QIAGEN Plasmid Plus Midi Kit.

**Lentiviral Production and Transduction**

Transduction protocols were adapted and optimised from published protocols[13].

**Lentiviral Packaging:** Lentiviral particles were generated using Lenti-X 293T cells. Transfection was performed using Lipofectamine 3000 according to the manufacturer's instructions. Packaging plasmids *psPAX2* and *pMD2.G* (addgene 12260 and 12259) were combined with the specific LentiGuide-Puro-P2A-EGFP sgRNA constructs (*Ptprj*, *Rpa3* and *Renilla*) in Opti-MEM medium. Following a 15 min complexation period, the transfection mix was added dropwise to the Lenti-X 293T cultures and incubated at 37°C overnight.

**Transduction (Spinoculation):** Supernatant containing lentiviral particles was harvested and centrifuged at 400 × *g* for 5 min before adding directly. Target $SpCas9\text{-}Cebpa^{N\text{-}term/N\text{-}term}$ and $SpCas9\text{-}Cebpa^{C\text{-}term/N\text{-}term}$ cells were counted and resuspended in complete culture medium supplemented with 20% FBS, IL-3, and polybrene (final concentration 2 µg/mL). Cells were seeded into 12-well plates at a density of 0.3-0.5 × $10^6$ cells/mL in 1.5 mL media. 1.5 mL viral supernatant was then added. Lentiviral transduction was performed via double spinoculation (900 × *g* for 90 min at 37°C) with a 5h incubation between spins. Before the second spin the plate is spun at 400 × *g* for 5 min, 1.5 mL is removed and replaced with more lentiviral supernatant. The following day the viral supernatant was removed, and cells were resuspended in fresh recovery medium (supplemented with 50 ng/mL mIL-3 and 20 ng/mL mIL-6) and transferred to 24-well plates for recovery and subsequent expansion.

**Validation of Transduction with Rpa3**

*SpCas9-Cebpa*$^{N\text{-}term/N\text{-}term}$ cells transduced with the *Rpa3* sgRNA were monitored for GFP by flowcytometry to confirm the sharp decrease in GFP+ cell number over time. Although GFP expression is variable quick disappearance compared with controls was observed.

***Ptprj* Sequencing in Patient Samples**

96 biallelic CEBPA-mutant AML samples were identified in the UCL/UCLH Haematology Biobank and DNA stored from associated clinical trials.

**Targeted Sequencing Design and Library Preparation:** A custom targeted sequencing panel for the PTPRJ gene was designed by Twist Bioscience (Design ID: TE-97414819). The panel comprises 236 probes covering 4,361 bp of the human genome (hg38) using a 4X tiling strategy. Genomic DNA libraries were prepared using the Twist Library Preparation Kit EF 2.0. Following enzymatic fragmentation (optimized for a target insert size of 375–450 bp) and end-repair, Twist Universal Adapters were ligated to the fragments, and libraries were amplified using Twist Unique Dual Indexed (UDI) Primers.

**Multiplexed Target Enrichment:** Samples were multiplexed in 8-plex pools. Each indexed library contributed 187.5 ng to the hybridization pool, resulting in a total input mass of 1,500 ng (1.5 µg) per hybridization reaction. Enrichment was performed using the Twist Target Enrichment Standard Hybridization v2 protocol, with libraries hybridized to the custom PTPRJ panel for 16 hours at 70°C. Captured targets were purified using streptavidin-coated magnetic beads and subjected to stringent washes. Enriched pools were amplified via post-hybridization PCR using 7 cycles, and final libraries were purified and validated for concentration and size distribution prior to sequencing on an Illumina platform.

**Sequencing and Read Processing:** Targeted next-generation sequencing of the *PTPRJ* locus was performed on a cohort of 96 samples utilizing pre-made libraries sequenced on an Illumina NovaSeq X Plus platform using a 25B flow cell with 150 bp paired-end reads. Raw sequencing data in FASTQ format were subjected to initial quality control using FastQC[14]. Low-quality bases, adapter sequences, and short reads were trimmed using Trimmomatic[15]. Raw sequencing reads were aligned to the human reference genome assembly GRCh38 using the Burrows-Wheeler Aligner (bwa-mem)[16]. Post-alignment processing, duplicate marking, standardizing sequence alignments in BAM format, and coordinate sorting and indexing were conducted via samtools[17] and picard[18]. Genomic regions of interest were constrained to a high-density target panel spanning the *PTPRJ* and *CEBPA* loci using a customized coordinate layout from Twist Bioscience.

**Variant Calling and Pipeline Validation:** To ensure robust identification of single nucleotide variants (SNVs) and small insertions/deletions (indels) within the target genomic regions of *PTPRJ*, a consensus variant calling strategy was implemented using three independent algorithms: BCFtools, MuTect2 (GATK), and VarScan2.

1. **BCFtools Pipeline[19]:** Raw variant calling was initially performed using the bcftools mpileup command to compute genotype likelihoods, followed by bcftools call (v1.19) utilizing the multiallelic calling model (-m).
2. **GATK MuTect2 Pipeline[20]:** Somatic and low-frequency germline variant detection was performed using MuTect2 within the Genome Analysis Toolkit (GATK, v4.4.0.0). The analysis was restricted to the targeted intervals using the validated *PTPRJ* BED file. To minimize common germline contamination, a maximum population allele frequency threshold was set. Raw MuTect2 calls were subsequently processed through GATK's FilterMutectCalls tool to separate true biological variants from sequencing and alignment artifacts based on internal error models.
3. **VarScan2 Pipeline[21]:** SAMtools mpileup was piped directly into the VarScan2 mpileup2snp tool to identify high-confidence single nucleotide polymorphisms (SNPs) in a single-sample stream, constrained by the target BED intervals. The algorithm was executed with default stringency filters, requiring a minimum sequencing coverage of 8×, a minimum of 2 variant-supporting reads.

**Variant Calling and Annotation:** To ensure high-confidence calls, a consensus-based approach was employed requiring variants to be detected by three independent variant calling algorithms. Variants were subjected to stringent quality filters, including a minimum read depth of 20x and a minimum support of 5 mutant reads per variant. Variants were annotated using the Variant Effect Predictor (VEP)[22]. Functional impacts were classified according to the Sequence Ontology standards. Mutation type (somatic vs. germline) was determined based on a Variant Allele Frequency (VAF) threshold of 30%[23], where values <30% were classified as somatic and values >30% were classified as germline.

**Supplementary Table 1**

Differential expression of colony stimulating factor receptors between *cebpa* mutants and wildtype, from RNA-Seq of HSC at 4dpf.

| | C-term/C-term | | N-term/N-term | | N-term/C-term | |
|---|---|---|---|---|---|---|
| **csf1ra**<br>M-CSFR<br>Macrophage | ns | NES = 0.66<br>p = 0.859 | ↓ | NES = -2.85<br>p = 0.036 | ns | NES = -0.74<br>p = 0.80 |
| **csf2rb**<br>GM-CSFR<br>Granulocyte-macrophage | ↑ | NES = 1.59<br>p = 0.0099 | ns | NES = 0.14<br>p = 0.87 | ns | NES = 0.03<br>p = 0.98 |
| **csf3r**<br>G-CSFR<br>Granulocyte | ns | NES = -1.50<br>p = 0.14 | ↓ | NES = -3.16<br>$p = 7.7x10^{-6}$ | ↓ | NES = -2.01<br>p = 0.017 |

**Supplementary Table 2**

Phosphoproteomics mass spectrometry: 28 MS2 scans of variable windows

| m/z | z | Isolation Window (m/z) |
|---|---|---|
| 409.6991 | 2 | 70.1 |
| 462.7067 | 2 | 38 |
| 495.2182 | 2 | 29.1 |
| 521.6214 | 2 | 25.8 |
| 545.6382 | 2 | 24.3 |
| 567.7783 | 2 | 22 |
| 588.7557 | 2 | 22 |
| 609.5119 | 2 | 21.6 |
| 630.3757 | 2 | 22.2 |
| 651.1157 | 2 | 21.3 |
| 671.1172 | 2 | 20.7 |
| 690.9056 | 2 | 20.9 |
| 710.3482 | 2 | 20 |
| 730.1012 | 2 | 21.5 |
| 750.5052 | 2 | 21.3 |
| 770.7238 | 2 | 21.1 |
| 792.1617 | 2 | 23.8 |
| 814.9678 | 2 | 23.9 |
| 837.7671 | 2 | 23.7 |
| 861.2793 | 2 | 25.3 |
| 886.4203 | 2 | 27 |
| 914.1665 | 2 | 30.5 |
| 944.5193 | 2 | 32.2 |
| 976.2617 | 2 | 33.3 |
| 1011.194 | 2 | 38.6 |
| 1051.426 | 2 | 43.9 |
| 1100.375 | 2 | 56 |
| 1163.425 | 2 | 72.1 |

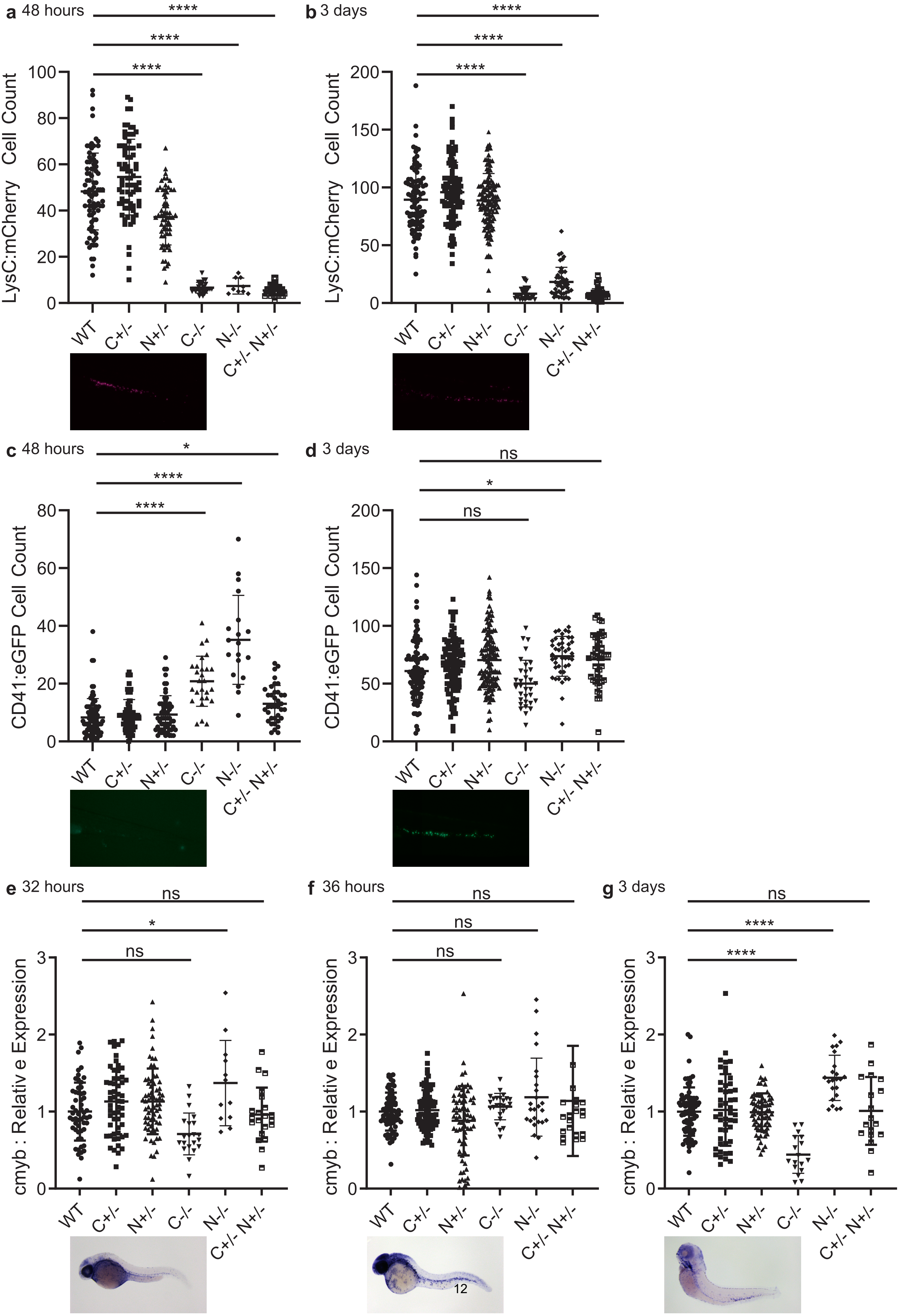

a 48 hours
b 3 days
c 48 hours
d 3 days
e 32 hours
f 36 hours
g 3 days
****
*
ns
LysC:mCherry Cell Count
CD41:eGFP Cell Count
cmyb : Relativ e Expression
WT
C+/-
N+/-
C-/-
N-/-
C+/- N+/-

**Supplementary Figure 1:** Automated cell counts showing *lysC:mCherry+* expression in the CHT of *cebpa* mutants at **a.** 48hpf and **b.** 3dpf. Automated cell counts showing *cd41:GFP*+ expression in the CHT of *cebpa* mutants at **c.** 48hpf and **d.** 3dpf. Normalised intensity *c-myb* expression by whole-mount in situ hybridisation in the CHT of *cebpa* mutants at **e.** 32hpf, **f.** 36hpf and **g.** 3dpf. One way ANOVA with GraphPad Prism. ns = $p > 0.05$; * = $p \leq 0.05$; ** = $p \leq 0.01$; *** = $p \leq 0.001$; **** = $p \leq 0.0001$.

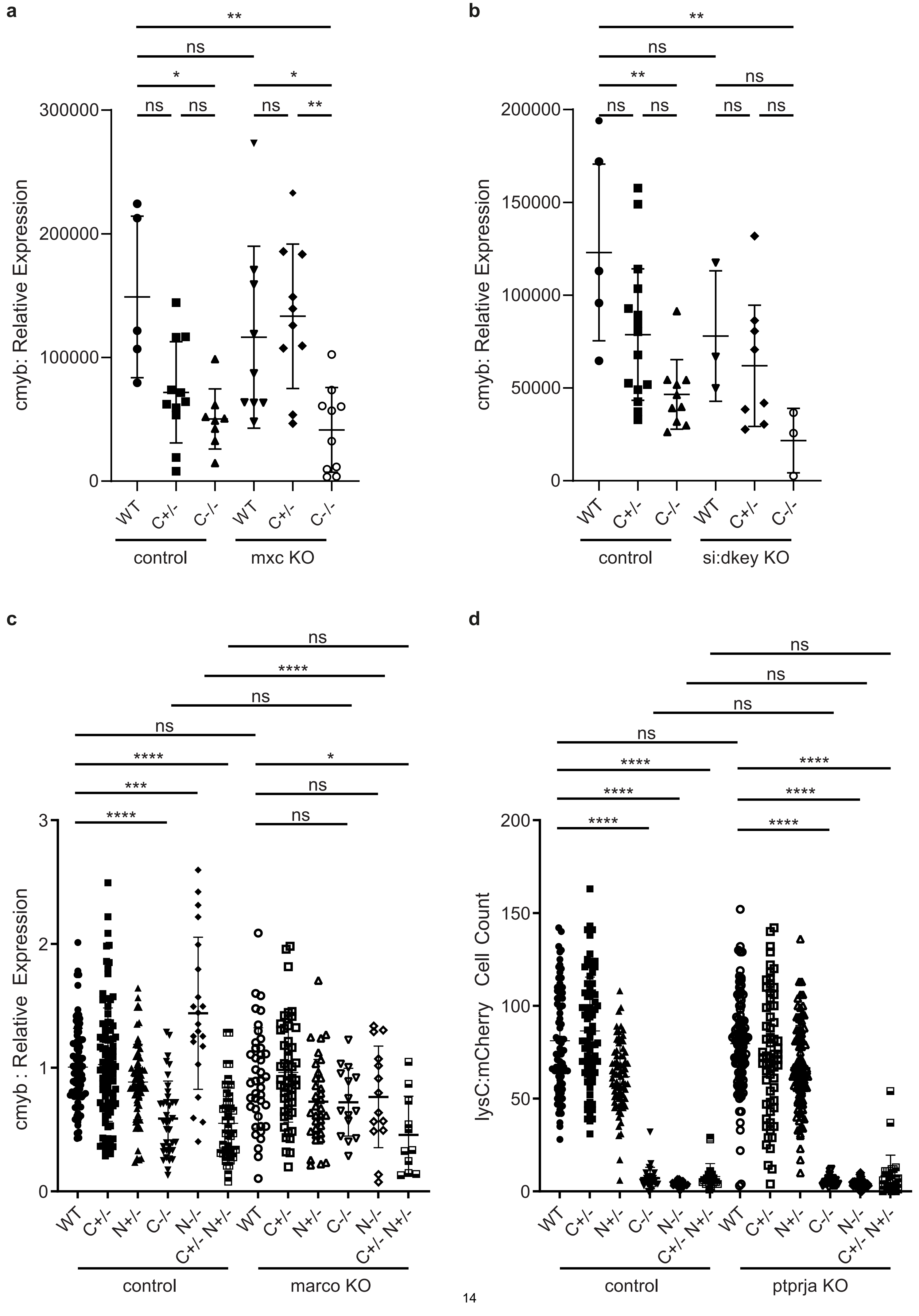

a
b
c
d
**
ns
*
ns
ns
ns
**
300000
200000
100000
0
cmyb: Relative Expression
WT
C+/-
C-/-
control
mxc KO
150000
50000
si:dkey KO
****
***
3
2
1
cmyb : Relative Expression
N+/-
N-/-
C+/- N+/-
marco KO
lysC:mCherry Cell Count
150
50
ptprja KO

**Supplementary Figure 2:** Normalised intensity *c-myb* expression by whole-mount in situ hybridisation in the CHT at 4dpf in: **a.** control and *mxc* (multi sex combs) CRISPR injected *cebpa*$^{C\text{-}term}$ mutants as part of the initial F0 screen, **b.** control and *si:dkey -148a17.5* CRISPR injected *cebpa*$^{C\text{-}term}$ mutants as part of the initial F0 screen, and **c.** control and *marco* CRISPR injected *cebpa* mutants. **d.** Automated cell counts showing *lysC:mCherry*+ expression in the CHT of *cebpa* mutants with and without loss of *ptprja* at 4dpf. Two way ANOVA with GraphPad Prism. ns = $p > 0.05$; * = $p \leq 0.05$; ** = $p \leq 0.01$; *** = $p \leq 0.001$; **** = $p \leq 0.0001$.

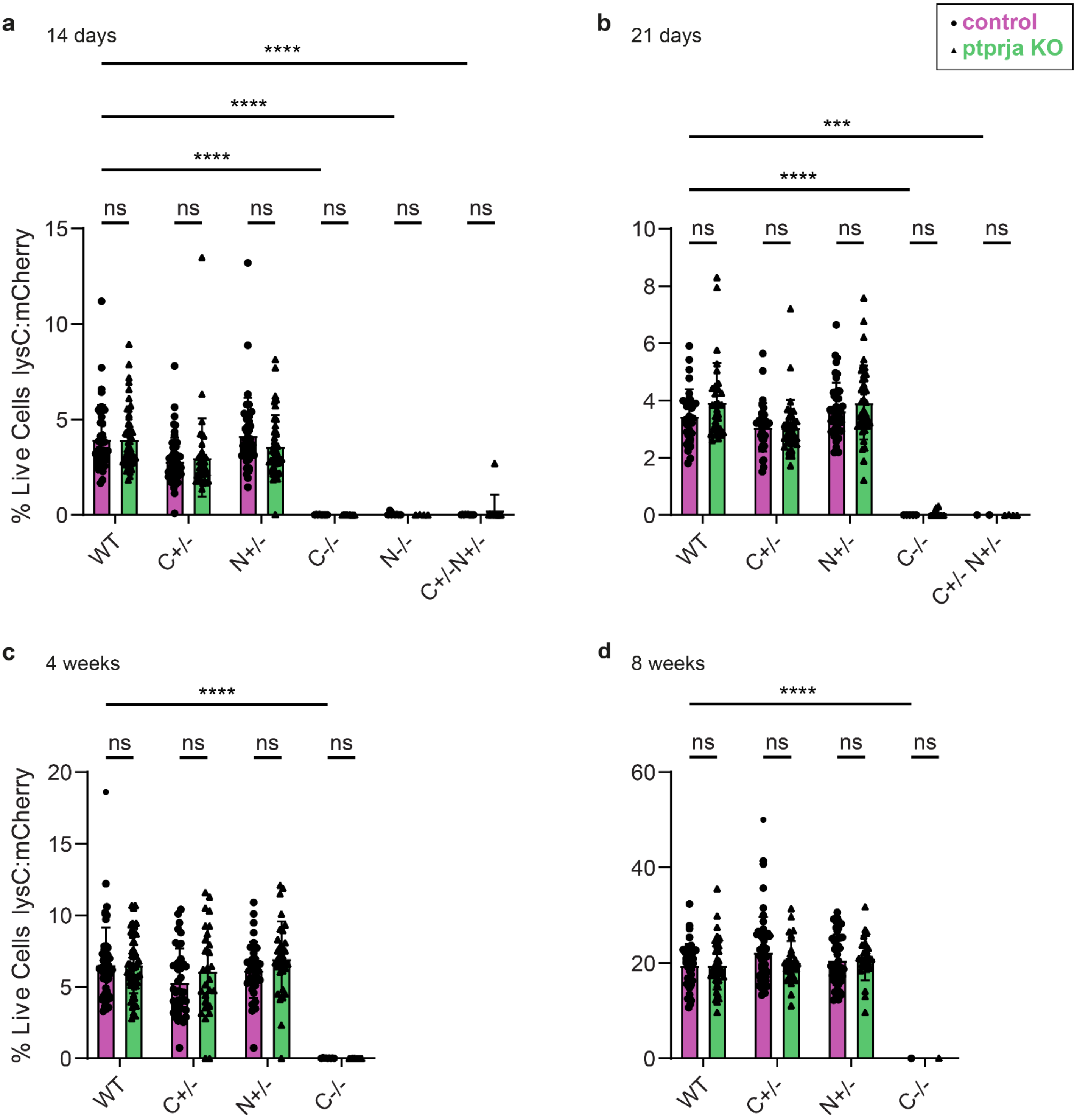


**Supplementary Figure 3: a.** Analysis by flow cytometry of *lysC:mCherry*+ HSC cells in control and *ptprja* CRISPR injected *cebpa* mutants at 14 days. **b.** Analysis by flow cytometry of *lysC:mCherry*+ HSC cells in control and *ptprja* CRISPR injected *cebpa* mutants at 21 days. **c.** Analysis by flow cytometry of *lysC:mCherry*+ HSC cells in control and *ptprja* CRISPR injected *cebpa* mutants at 4 weeks. **d.** Analysis by flow cytometry of *lysC:mCherry*+ HSC cells in control and *ptprja* CRISPR injected *cebpa* mutants at 8 weeks. All biallelic *cebpa* mutants have an absence of myeloid cells at all ages.

**a**

| Patient ID | Location | VAF | Nucleotide | Amino Acid | Mutation Type | Consequence | Impact |
|---|---|---|---|---|---|---|---|
| B_A01 | 11_47980983_C | 0.14% | T>C | Leu24Pro | somatic | missense | Moderate |
| B_A04 | 11_47980983_C | 0.13% | T>C | Leu24Pro | somatic | missense | Moderate |
| B_A09 | 11_47980983_C | 0.28% | T>C | Leu24Pro | somatic | missense | Moderate |
| B_B01 | 11_48137137_A | 0.42% | G>A | Ala670Thr | somatic | missense | Moderate |
| B_B02 | 11_48137137_A | 0.25% | G>A | Ala670Thr | somatic | missense | Moderate |
| B_B08 | 11_47980983_C | 0.40% | T>C | Leu24Pro | somatic | missense | Moderate |
| B_B08 | 11_48137137_A | 0.55% | G>A | Ala670Thr | somatic | missense | Moderate |
| B_B12 | 11_48127960_A | 0.52% | G>A | Arg425His | somatic | missense | Moderate |
| B_G09 | 11_48127960_A | 37.90% | G>A | Arg425His | germline | missense | Moderate |

**b**

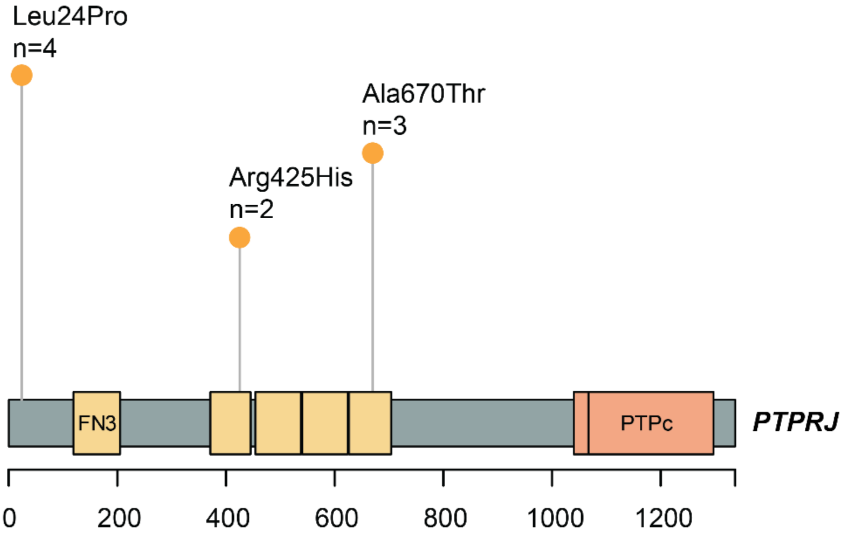


**Supplementary Figure 4:** 96 human biallelic *CEBPA*-mutant AML samples were sequenced for mutations in *PTPRJ*. **a.** High confidence variants independently called by three independent algorithms: BCFtools, MuTect2 (GATK), and VarScan2. Variants were annotated using the Variant Effect Predictor (VEP). **b.** Lollipop plot showing location of the mutations in the *PTPRJ* gene.
